# Characterizing the Prey Landscape of Endangered Cook Inlet Beluga Whales Using an eDNA Time Series

**DOI:** 10.64898/2026.11.12.766312

**Authors:** Zachary Gold, Andrew Olaf Shelton, Deborah D. Boege Tobin, Ana Ramón-Laca, Sonia V Kumar, Madison M. Kosma, Teresa Becher, Verena Gill, Paul Wade, Kim M. Parsons

## Abstract

Our ability to understand habitat utilization in the context of foraging, prey dynamics, and anthropogenic disturbance is critical for the effective management of protected species globally. Here we examine the occurrence of the endangered Cook Inlet beluga whale (*Delphinapterus leucas*) in the context of seasonally changing prey fields by leveraging quantitative eDNA metabarcoding approaches. Specifically, eDNA approaches were used to generate estimates of fish abundances using a joint modeling framework that derives total DNA concentration from a coho (*Oncorhynchus kisutch*) qPCR assay and bias-corrected proportional abundance of multiple salmonid species using MiFish *12S* metabarcoding of environmental and mock community samples. Temporal patterns of beluga whale occurrence were documented in two estuarine systems using visual observations, metabarcoding, and a novel beluga whale qPCR assay. We demonstrate that weekly time series before, during, and after beluga whale occurrence provide sufficient resolution to resolve changes in predator occurrence and prey field shifts across Cook Inlet. Specifically, we observed Cook Inlet beluga whales co-occurring with increases in coho DNA concentrations in the vicinity of Twentymile River at the terminus of Turnagain Arm. Monthly sampling on the Kenai River captured known seasonal anadromous fish migration patterns and identified a clear mismatch between Cook Inlet beluga whale habitat utilization and spring eulachon (*Thaleichthys pacificus*) runs. Our results also provide valuable insights to improve quantitative metabarcoding approaches through informed selection of qPCR targets and mock communities to support future eDNA-based studies in the highly turbid, dynamic habitats of Cook Inlet. Specifically, our results highlight the need for increased temporal sampling frequency as opposed to additional spatial and depth sampling in systems with extreme tides, as well as the use of larger pore sizes (> 5µm) to facilitate filtration in samples with high suspended glacial silt loads. We also document the capacity for eDNA signatures to integrate across 18 km spatial scales during exceptionally large tidal bores (> 10 m) that characterize Turnagain Arm. Ultimately, this study demonstrates the value of pairing visual surveys of endangered Cook Inlet beluga whales with strategic shore-based eDNA sampling to generate information on the co-occurrence of Cook Inlet beluga whales and putative prey fields.

## Introduction

Marine mammals are culturally, economically, and ecologically important species globally. However, many species are under threat from a variety of anthropogenic impacts including overharvesting, biomagnification, acoustic pollution, and loss of foraging habitat and prey abundance (Avila et al., 2018; Booth et al., 2020; Duarte et al., 2021; Erbe et al., 2018; Gulland et al., 2022; Harkonen et al., 2012; Nelms et al., 2021; Pompa et al., 2011). Often, multiple stressors impact such threatened marine mammal populations, leading to cumulative effects on a population, making it difficult to identify the relative impacts of concomitant stressors and limiting our ability to implement effective management and conservation actions (Maxwell et al., 2013; McHuron et al., 2023; Murray et al., 2021; National Academies of Sciences, Engineering, and Medicine et al., 2017; Williams et al., 2016).

Cook Inlet beluga whales (*Delphinapterus leucas*) are a small, endangered population of resident whales impacted by multiple anthropogenic factors including physical and acoustic disturbance, chemical and biological pollution, and prey limitations (Castellote et al., 2019; McHuron et al., 2023; Warlick et al., 2024). This isolated, genetically distinct population segment (DPS) of resident beluga whales resides exclusively in Cook Inlet (Figure 1), seasonally migrating within the 20,000 km^2^ inland sea (O’Corry-Crowe et al., 2018, 2020). However, over a decade after the federal listing of this endangered DPS under the U.S. Endangered Species Act, the lack of recovery of Cook Inlet beluga whales remain a significant management challenge. The Cook Inlet beluga whale DPS was estimated to be 1,300 in the late 1970s (Calkins, 1989), and was reduced due to unsustainable levels of harvest (Hobbs et al., 2000; Mahoney and Shelden, 2000). As of 2023, the ‘best’ estimate of abundance for the population based on aerial survey data is 331 whales (95% probability interval 290 to 386) with a largely stable population since 2004 (Goetz et al., 2023).

**Figure 1.**
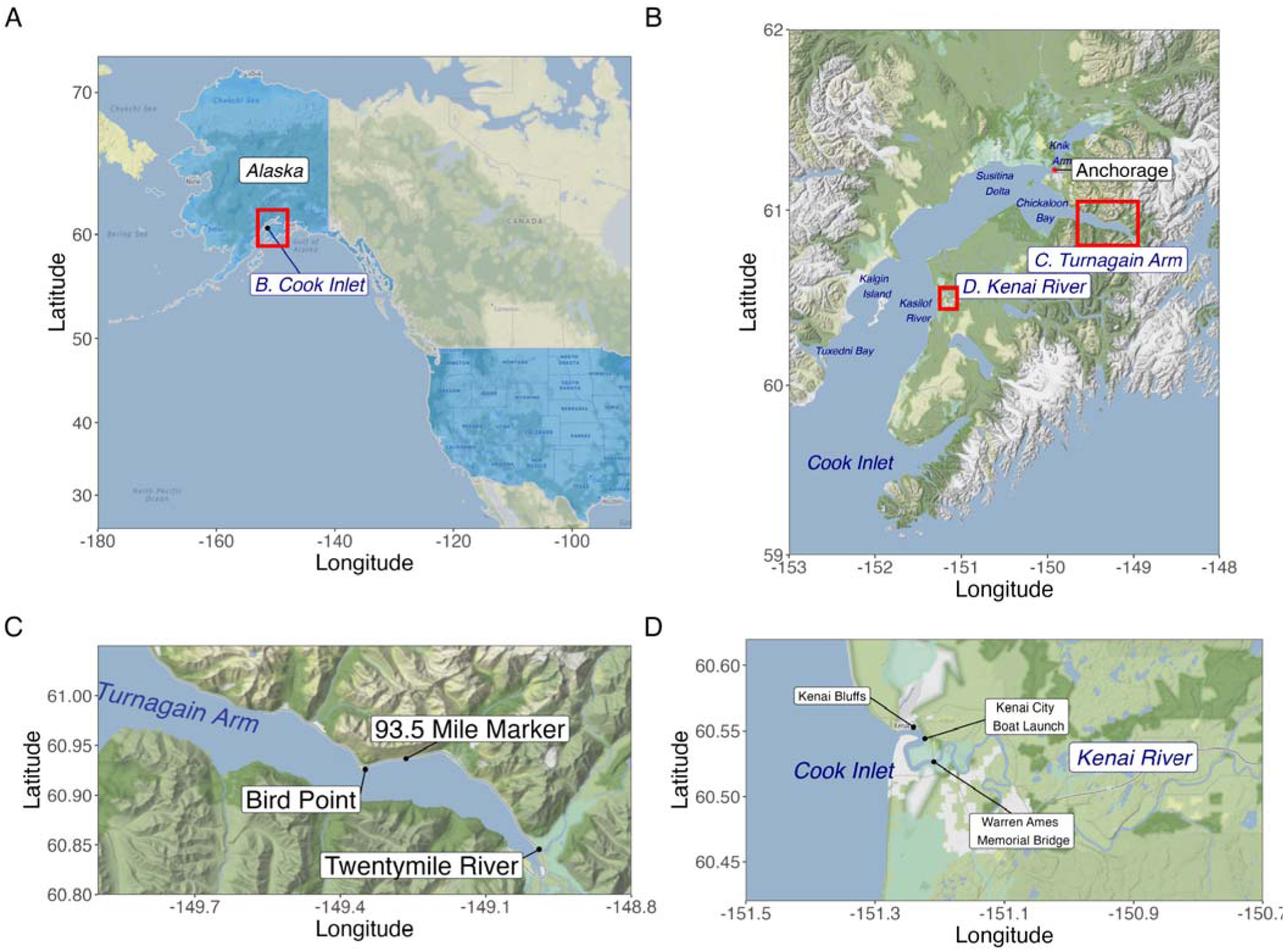
Map of Alaska (A), Cook Inlet (B) with Turnagain Arm (C) and Kenai River (D) sites. On the Kenai River, visual surveys were conducted at Kenai Bluffs and Kenai City Docks while eDNA samples were collected at Kenai City Boat Launch and Warren Ames Memorial Bridge. In Turnagain Arm, visual surveys were conducted at 95.3 Mile Marker and the mouth of the Twentymile River while eDNA samples were collected at Twentymile River.

The three threats of high relative concern to Cook Inlet beluga whale recovery are catastrophic events (e.g., natural disasters; oil spills; mass strandings); cumulative effects of multiple stressors; and noise (Recovery Plan for the Cook Inlet Beluga Whale (Delphinapterus Leucas), n.d.; Shelden et al., 2021). In particular, synergistic effects between anthropogenic activities and prey having the potential to further reduce suitable foraging habitat (Castellote et al., 2019). Anthropogenic impacts on beluga whale prey availability and habitat quality are particularly concerning in light of an observed range contraction of Cook Inlet beluga whales into the upper Cook Inlet in the summer, which is also home to over 40% of Alaska’s human population and significant vessel, aircraft, marine construction, seismic survey, and military activities (Castellote et al., 2019; D. J. Rugh et al., 2010). In addition, during the past decade, the abundance of Chinook (*Oncorhynchus tshawytscha*) salmon populations and forage fishes within the region, known Cook Inlet beluga whale prey species (Quakenbush et al., 2015), have declined (Arimitsu et al., 2021; Castellote et al., 2021; Jones et al., 2020; McHuron et al., 2023; Oke et al., 2020; Suryan et al., 2021), emphasizing the need to better understand the interplay among anthropogenic noise, Cook Inlet beluga whale foraging behavior, and prey abundances to inform conservation efforts.

Long term monitoring of whales from visual and acoustic observations has provided important insights into seasonal habitat utilization within Cook Inlet (Brewer et al., 2023; Castellote et al., 2020, 2021; Goetz et al., 2023; McGuire, Stephens, et al., 2020; O’Corry-Crowe et al., 2018). Cook Inlet beluga whales have been routinely observed in Turnagain Arm, Chickaloon Bay, Susitina Delta, Knik Arm, Kenai and Kasilof Rivers, Tuxedni Bay, and Kalgin Island (Figure 1) during the spring through summer season, moving among locations, often exhibiting predictable seasonal movements (Castellote et al., 2020; Goetz et al., 2023; McGuire, Stephens, et al., 2020; McHuron et al., 2023). Interestingly, whales appeared to move between sites in the Susitna Delta and Chickaloon Bay during summer months and transition to Knik Arm and Turnagain Arm in the fall. Shore-based observers routinely document the return of beluga whales to Twentymile river in Turnagain Arm every spring (May-June) and fall (August-November) noting a general absence from the areas in the summer months (https://akbmp.org/; (Goetz et al., 2023). Likewise, Cook Inlet beluga whales have been routinely observed in spring and fall in Kenai River with a notable absence in summer months during the sockeye run in recent years (https://akbmp.org/ and (Kumar et al., 2024) in contrast to historical observations (Huntington, 2002). Together, these observations demonstrate strong seasonal associations with the riverine and estuarine systems within Cook Inlet, likely driven by changes in the distribution and availability of key fish prey species (Calkins, 1989; Castellote et al., 2020; Hobbs et al., 2005; Huntington, 2002; D. Rugh et al., 2002; Shelden et al., 2022; Wolf et al., 2018).

Unfortunately, obtaining prey consumption data for Cook Inlet beluga whales is extremely challenging given the highly turbid and tidally dynamic system which precludes the collection of fecal samples or underwater observations (Castellote et al., 2020; McHuron et al., 2023). Existing diet data comes from the analysis of prey remains from a limited number (n=18) of stomachs collected between 2002 and 2012 from harvested or stranded whales (Quakenbush et al., 2015). Similar to other Alaska beluga whales, their diet was dominated by anadromous fishes, particularly from spring through fall (Castellote et al., 2021; Quakenbush et al., 2015) including salmonids [coho (*Oncorhynchus kisutch*), Chinook (*O. tshawytscha*), and chum (*O. keta*)] and smelt [eulachon (*Thaleichthys pacificus*)]. Other Alaska beluga whale populations are known to consume anadromous fish species not observed in the stomachs collected from Cook Inlet beluga whales, including sockeye (*O. nerka*) and pink salmon (*O. gorbuscha*) in Bristol Bay as well as rainbow smelt (*Osmerus mordax*) and capelin (*Malloutus villosus*) in the Eastern Bering sea (Quakenbush et al., 2015). Such work aligns with traditional ecological knowledge of Cook Inlet beluga whales utilizing Kenai River during summer sockeye runs (Huntington, 2002) as well as Turnagain Arm during fall coho runs (Paul Crews *pers. comm,*). Cook Inlet beluga whale stomach contents contained gadids (saffron cod (*Eleginus gracilis*), walleye pollock (*Gadus chalcogrammus*), Pacific cod (*Gadus macrocephalus*)) which are known to frequent shallow waters and estuaries of Cook Inlet during juvenile life stages (Guo et al., 2022; Quakenbush et al., 2015). Information gained through the analysis of stomach contents provides critical baseline data on beluga whale prey preferences. However the small sample size and unknown provenance of some samples limits population level inference, emphasizing the need for alternative approaches.

Comparing temporal patterns of co-occurrence for predators and prey can offer valuable insights on targeted prey species (Carroll et al., 2019), especially when predator and prey exhibit correlated changes in abundance and clear signs of foraging behavior (Amir et al., 2022; Blanchet et al., 2020; Ford & Goheen, 2015). Traditionally, acquiring such data in Cook Inlet is difficult given the highly dynamic tidal environment (tides > 10 m; with large tidal bores; (Fissel & Lin, 2022; Oey et al., 2007; Wang & Yang, 2020)) and the lack of biomonitoring efforts for fish species without high commercial value, with only a few salmon runs monitored, such as on the Susitna River. Thus, there is a need for novel, easy-to-deploy approaches to successfully characterize the prey landscape and link these observations to beluga whale occurrence. Developing such spatially and temporally resolved putative prey targets across Cook Inlet can better inform Cook Inlet beluga whale habitat utilization patterns within the region and inform critical habitat needs (McHuron et al., 2023).

To address this challenge, we used quantitative environmental DNA (eDNA) approaches to characterize prey dynamics in two key riverine systems used by Cook Inlet beluga whales and related these patterns to beluga whale occurrence from visual observations and eDNA detections. In conjunction with the Alaska Beluga Monitoring Program (AKBMP), a NOAA Fisheries-led citizen science program, we conducted time-series eDNA sampling in the Twentymile River during the seasonal beluga-use period and in the Kenai River across an annual cycle to characterize the prey landscape associated with beluga whale occurrence. Our results show that quantitative eDNA approaches can reveal shifts in predator occurrence and prey fields, providing new insight into the putative seasonal foraging patterns of this endangered population.

## Methods

We present a summarized description of the methods here with a full detailed description in the Supplemental Materials Appendix 1. All sequences are made available through GenBank SRA (XX), and data and code through Zenodo (10.5281/zenodo.24183095). The possession, collection and processing of cetacean tissues and eDNA samples was conducted under NOAA’s Marine Mammal Laboratory research permit (#20465).

### Visual Beluga Whale Observations

Visual observations of beluga whale were conducted by co-authors or AKBMP volunteers following the AKBMP monitoring protocol (https://akbmp.org/resources/). Monitoring session timing was based on NOAA-predicted high tides, using the Anchorage, Knik Arm station (NOAA station 9455920) for Twentymile River and the Kenai River Entrance station (NOAA station TWC1983) for Kenai River. Monitoring sessions lasted two hours, beginning at Anchorage high tide in Twentymile River and five hours before high tide in the Kenai River (Figure 1). We compared eDNA detections of beluga whales at Twentymile River to the spatio-temporally concurrent visual observations at both Twentymile river (60.8458 N, 148.9903 W) and Highway Mile Marker 95.3 Pullout (60.9324739°N, 149.3234756°W), approximately 18 km downstream in Turnagain Arm. In 2021, AKBMP monitoring at Twentymile River began on August 4 and ended on November 12 while monitoring at Mile Marker 95.3 Pullout began August 15 and ended on November 15. We note that *ad hoc* visual observations from AKBMP volunteers began at Mile Marker 95.3 and Bird Point on August 4th, 2021 which we use solely as positive confirmation of Cook Inlet beluga occurrences (S. Steinhert *pers. comms*).

On the Kenai River, we compared visual observations made at either the Kenai Bluffs (60.552895°N, 151.240538°W) or City of Kenai Boat Launch (60.544138°N, 151.222841°W) to our two Kenai River eDNA sampling locations, approximately 1.7 km from the City of Kenai Boat Launch (60.544138°N, 151.222841°W) and 5.56 km at the Warren Ames Memorial Bridge. eDNA occurred from November 2020 to November 2021. Visual observations on the Kenai River we conducted from April 2021 to October 2021.

### eDNA Sampling

#### Twentymile River

In 2021, we conducted sub-monthly eDNA sampling on Twentymile River at the same location as visual observations to characterize both fish prey community composition and beluga whale occurrence. Specifically, we collected three eDNA samples three to four times per week at the surface (0 m) from August 1 to August 26, followed by bi-weekly sampling until November 11 (Figure 1, Table S1) coinciding with peak high tide. Samples were collected using a Nasco Swing Sampler (Fisher Scientific Catalogue No. 01-812-84) with a one liter Nalgene bottle mount. Bottles were sterilized in the lab with 10% commercial bleach for one hour between uses and triple rinsed with DI water to remove all traces of bleach. Prior to sample collection events, bottles were rinsed in the river three times at the sampling location. Sealed and labeled Nalgene bottles were stored in a cooler on ice before being transported to Girdwood, Alaska for filtering. Samples were vacuum filtered (see below) within two hours of sample collection (Table S2).

#### Kenai River

We conducted monthly eDNA sampling on the Kenai River at two sites, the City of Kenai Boat Launch, near the mouth of the river, and Warren Ames Memorial Bridge, 5.56 km upstream (Figure 1). eDNA samples were collected monthly at both locations, at both the surface (0 m) and five meter depth from November 14th, 2020 to November 14th, 2021 (Table S1). Samples were collected using a weighted two L Kemmerer attached to a rope with marked lengths, deployed over the side of the railing. At each site and depth, three replicate one liter samples were collected at each time point. Approximately 500 mL of water was used from each Kemmerer to rinse one liter Nalgene bottles before sampling. Samples were then stored on ice in coolers before being transported to the Kachemak Bay Campus of Kenai Peninsula College-University of Alaska Anchorage, Homer, Alaska for filtering. Samples were collected at high tide or as close to high tide as feasible. Given logistical constraints, samples were often transferred to refrigerators for overnight storage prior to being filtered the following day (Table S1).

### eDNA Filtration & Preservation

Each eDNA sample consisted of one liter of water vacuum filtered following Ramón-Laca et al., 2021 using single use 250 mL Millipore Stericups (Sigma Aldrich) fitted with 1 µm 47mm Nitrocellulose Mixed Ester (MCE) Sterlitech filters. Vacuum pressure was applied until the entire one liter sample was completely filtered (maximum time was 3 hours 19 minutes due to high glacial sediment loads). Filters were then transferred with sterile forceps to either 5 mL tubes with 4 mL of Longmire lysis buffer (Kenai samples) or 1.5 mL tubes with 1 mL Longmire lysis buffer (Twentymile River) and stored at room temperature until DNA extractions were performed at the Northwest Fisheries Science Center in Seattle, WA.

### eDNA Extraction

eDNA was extracted from preserved filters using a modified Qiagen DNeasy Power Soil Pro Kit and a Qiacube liquid handling robot. Specifically, samples were first centrifuged before transferring the filter to labeled PowerBead Pro Tubes using a 1000 µL pipette tip and sterile flat tip forceps. The remaining Longmire’s lysis buffer was centrifuged at 6,000 g for 30 minutes at room temperature to pellet any lysed DNA. The supernatant was then removed and 800 µL of CD1 buffer was added to pellet, mixed 10 times using a micropipette, and transferred to the PowerBead Pro Tubes containing the filter. The rest of the extraction followed the manufacturer’s instructions on a Qiagen QIAcube automated DNA extraction instrument.

### eDNA Metabarcoding

Metabarcoding was performed on all eDNA samples using a two-step PCR library preparation method (Appendix 1). First, samples were amplified using the MiFish Universal Teleost primers (Miya et al., 2015) modified with Illumina Nextera adapter sequences following Curd et al., 2019.

We used the Anacapa Toolkit (Curd et al., 2019) for amplicon sequence variant (ASV) parsing, quality control, and taxonomic assignment (Curd et al., 2019; Gold et al., 2021) and a custom R script was used for post processing to conduct quality control on samples, ASVs, and species (see Appendix 2 for full description). We note that *Oncorhynchus* salmonid taxonomic assignments were reassigned using a custom script to deal with the high similarity in sequence alignments between ASVs. We first removed one reference barcode sequence of chum salmon (*O. keta*) deposited on GenBank (Accession LC091629.1, UW 042256) from Lake Pernatoye, Russia in 1997 with greater similarity to three other *O. nerka* accessions (LC835958.1, OL806600.1, LC340053.1) and appears to be either misidentified in the field, mishandled in the lab, or evidence of hybridization that is not relevant to species identification in Alaska. We then used the *DECIPHER* R package function *clusterize* to cluster both ASVs and reference sequences into bins at most 1 bp different from each other (Wright, 2016, 2024). We then accepted the taxonomic assignment provided by *Anacapa Toolkit* for ASVs and reference sequences that contained the same species level identification within a cluster. For clusters with conflicting species level identifications, ASVs were run through blast against NCBI GenBank. Species level taxonomic assignments were then only retained if all best matches (same percent identity) were to the same species, otherwise we assigned ASVs to the genus level (*Oncorhynchus* spp.).

### Species-specific qPCR assays

We ran four species specific qPCR assays on all eDNA samples targeting beluga whale, Chinook, coho, and eulachon. The Chinook and coho COIII/ND3 assays (Shelton et al., 2019) were multiplexed with a eulachon COI assay (Ramón-Laca et al., 2021) see Appendix 3 for multiplex assay validation). We also employed a new *12S* beluga whale specific qPCR assay (JUN). To create this assay, we pulled down all publicly available beluga whale mitochondrial sequences for all geographic regions, as well as other outgroup cetacean species, into Geneious (Dotmatics) and identified a species specific region of the hypervariable *12S* locus (see Appendix 3 for beluga whale assay design and validation).

All eDNA samples and non-template controls were run in triplicate in 10 μl reactions containing 1 × TaqPath ProAmp Multiplex Master Mix (ThermoFisher Scientific, Waltham, MA USA), forward and reverse primers (0.9 μM), hydrolysis probes (0.2 μM), and 2 μl of genomic DNA standard or eDNA. Cycling conditions were: initial denaturation step at 95°C for 10 min, followed by 45 cycles of 95°C for 15 s and 60°C for 1 min. Both assays included a synthetic internal positive control (IPC) to test for inhibition (*Euryapteryx curtus;* (Ramón-Laca et al., 2021) and at least six non template controls.

We generated a standard curve for each run containing equal concentrations of both target and IPC using a serial dilution of synthetic, linear, double-stranded oligonucleotide gBlocks^®^ Gene Fragments, herein ‘gBlocks’ (Integrated DNA Technologies, Coralville, IA, US): 100,000 copies µL^-1^, 10,000 copies µL^-1^, 1,000 copies µL^-1^, 100 copies µL^-1^, 10 copies µL^-1^, 5 copies µL^-1^, and 1 copy µL^-1^. All real-time PCRs were performed on an Applied Biosystems QuantStudio 6 Flex Real-Time system (ThermoFisher Scientific, Waltham, MA, US).

We processed raw qPCR data following the methods of Allan et al., 2023 and Guri et al., 2026 which built on similar linear models described in (McCall et al., 2014) and (Shelton et al., 2019). Here, the linear model accounts for nondetections as a function of DNA concentration within a sample with greater uncertainty for samples with lower concentrations, more realistically relating environmental samples to the standard curve. The linear model was fitted independently to each plate specific standard curve to estimate DNA concentrations from unknown environmental samples. qPCR data was processed in R using Stan (Carpenter et al., 2017) to employ the Allan et al. 2023 model (see Appendix 13). Samples with observed high standard deviations (coefficient of variation > 1.5) across technical replicates (distinct, independently run qPCR reactions using the source template DNA) were removed from the dataset. Samples with a Ct less than 40 cycles were considered positive detections.

### Quantitative Salmon Metabarcoding

We followed previously published methods to generate quantitative metabarcoding data from the MiFish sequence data to move beyond relative proportions of detected species (Allan et al., 2023; Guri et al., 2026). Specifically this model uses absolute abundance estimates derived from species-specific qPCR assays to anchor proportional estimates of multiple species derived from bias corrected metabarcoding data. DNA concentrations were not adjusted for environmental flow rates due to a lack of reliable accurate information on tide gauge data for Twentymile River (Figure 2).

**Figure 2.**
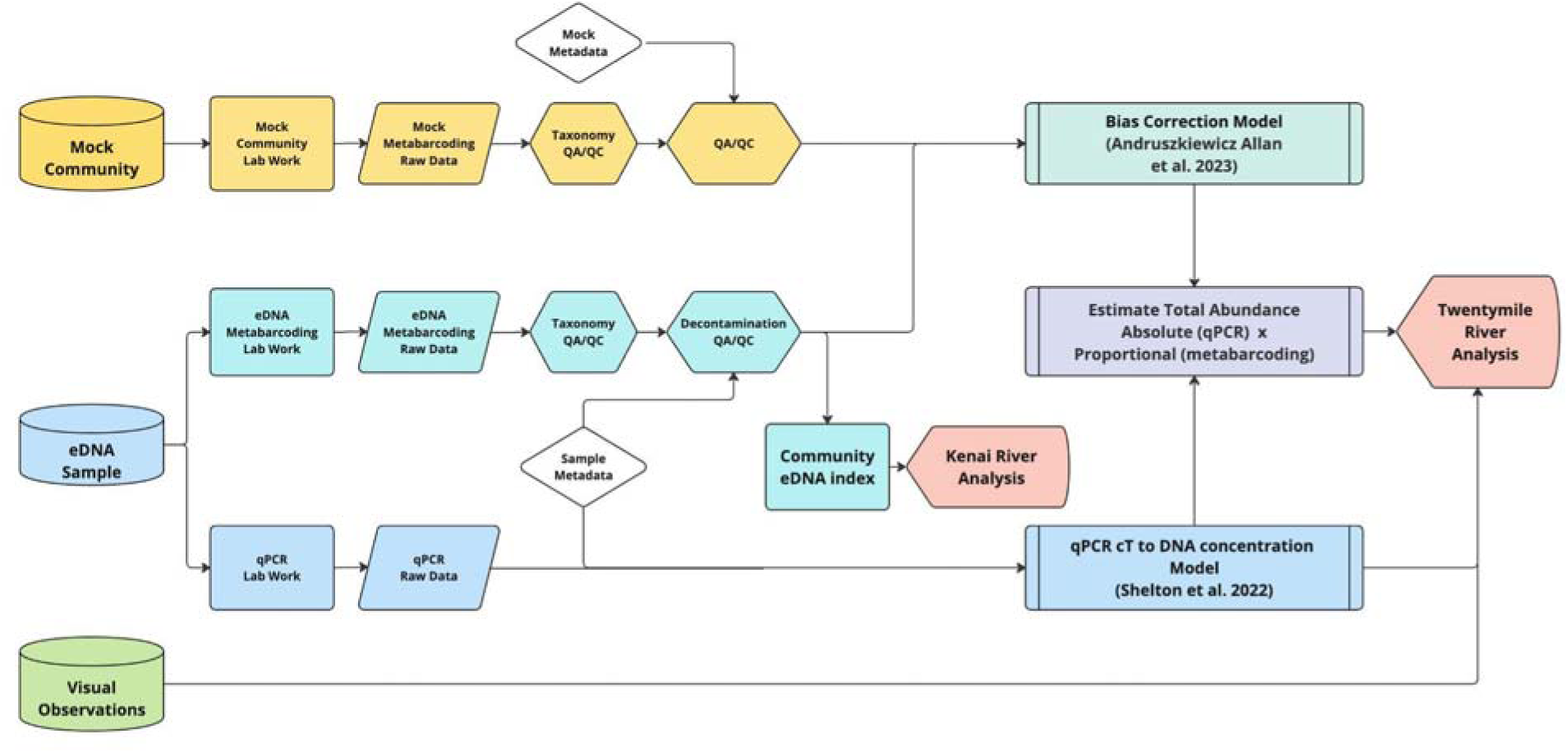
Quantitative Metabarcoding Workflow Diagram. Both metabarcoding and qPCR data were generated from all Cook Inlet eDNA samples. Raw metabarcoding data generated biased proportional abundance information due to differential amplification efficiencies. Species-specific amplification biases were identified from mock community metabarcoding data and subsequently used to correct amplification bias in the compositional eDNA metabarcoding data. Species-specific qPCR assays generated absolute abundance estimates for coho salmon in eDNA samples. Estimates of total abundance were calculated by multiplying biased corrected proportions from metabarcoding data by absolute DNA concentrations from qPCR. Results from the Kenai River strongly suggest swamping from metabarcoding data and thus quantitative metabarcoding approaches were not applied.

We utilized mock communities constructed from vouchered DNA specimens described in (Shelton et al., 2023) and (Gold et al., 2023) to estimate and correct for the known effects of species-specific PCR amplification bias (Shaffer et al., 2025). The highest amplification efficiency in the mock community was exhibited by Chinook followed by sockeye, coho, pink, chum, and steelhead (Appendix 4). Observed species-specific amplification efficiencies were used to calibrate the metabarcoding data to provide more accurate proportions of DNA in environmental samples. We note that we did not derive quantitative estimates for other species observed in metabarcoding data as we lacked accurate amplification efficiency estimates from mock community data.

We anchored the quantitative metabarcoding results to coho (*O. kisutch*) qPCR DNA concentrations, estimating the total amount of DNA in each sample by dividing the qPCR derived absolute abundance estimates of coho by the estimated metabarcoding derived proportional abundance of coho. We then estimated the absolute abundance of other salmon species by multiplying the metabarcoding derived proportional estimates of each species against the estimated total absolute DNA concentration in each sample. Coho was chosen as the anchor for total DNA concentration estimates because of its prevalence and abundance across environmental samples (mean 23,858 copies per L and >13,670 read abundance per sampling event). See Appendix 5 for comparisons using Chinook to anchor total DNA concentration estimates.

## Analyses

### Detecting Cook Inlet Beluga Whale with eDNA

We defined temporal periods of beluga whale occupancy in Turnagain Arm and Kenai River using visual observations collected during this study. Here we define the period prior to the first beluga observation as the period “before” beluga whale occupancy, and define the period between the first and last visual observation as the period “during” beluga whale occupancy. The occupancy period in Turnagain Arm was defined using visual observation data from both Mile Marker 95.3 Pullout and Twentymile River (Figure 1). In the Kenai River, we similarly defined the period of beluga whale occupancy as the period between the first and last visual observation for both the spring and fall seasons. We then compared beluga whale detection rates, here defined as the number of detections divided by the number of detection opportunities (e.g., a single sampling event inclusive of all replicates), across visual, qPCR, and metabarcoding datasets both during and outside of periods of beluga whale occupancy.

### Quantitative Metabarcoding Reveals Co-Occurrence of Beluga Whale and Salmon Prey in the vicinity of Twentymile River

To identify putative prey targets, we compared the total estimated DNA concentration of sockeye (*O. nerka*), coho (*O. kisutch)*, pink (*O. gorbuscha*), chum (*O. keta*), and Steelhead (*O. mykiss*) before and during Cook Inlet beluga whale occupancy in Turnagain Arm. We specifically assumed that the concentration of DNA was reflective of the underlying abundance of each species (Tillotson et al., 2019; Allan et al., 2023; and see Discussion). Here we focussed on changes in salmonid abundance because 1) we had a limited diversity of our mock community, 2) salmonids comprised 83.1% of total observed reads, 3) salmonids are a key prey source in the summer for Cook Inlet beluga whale (Quakenbush et al., 2015), and 4) we observed few eulachon during this time period (see Discussion).

### Spatial and Temporal Variation of eDNA Signatures on Kenai River

Using eDNA samples from the Kenai River, we compared changes in fish assemblages across season, site, depth, and biological replicates (here defined as distinct replicate water bottles) to inform future sampling efforts and the relative importance of spatial, temporal, and depth variation across eDNA sampling in a highly dynamic tidal system. Specifically, we transformed observed eDNA communities by first calculating percent abundances and then calculating Aitchison distances between each sample using *microviz (Barnett et al., 2021)*. We then ran a PERMANOVA on these centered log-ratio transformed communities and compared the R^2^ of season, site, depth, biological replicate, and their interactive effects (Dixon, 2003; McMurdie & Holmes, 2013).We had low statistical power to compare changes in putative prey abundance before, during, or after Cook Inlet beluga whale occupancy on the Kenai River because of the occurrence of molecular “swamping” (see Appendix 7) and infrequent, monthly sampling intensity. Therefore, we did not evaluate co-occurrence of Cook Inlet beluga whales and salmonid prey abundance in the Kenai River.

## Results

We collected a total of 258 eDNA water samples from 3 sites across two rivers spanning 12 months (Table S1). We generated 364 metabarcoding PCR technical replicates resulting in 52.0 million raw sequence reads across five MiSeq runs. After processing via the Anacapa Toolkit and subsequent decontamination, we retained results from 190 eDNA samples and 256 PCR technical replicates, yielding a total of 15,920,866 MiFish Teleost reads representing 64 species including 30 fish, 18 mammals, 13 birds, 2 sharks, and 1 lamprey (Table S2). Sample read depth ranged from 19,283 to 220,454 reads (mean 64,961 reads). Discarded samples and PCR replicates were removed because of low read depth (20,000 reads) or a single species comprising over 90% of captured reads.

From the same eDNA samples, we generated qPCR data from 4 species-specific assays with triplicate technical PCRs. We retained reliable concentration data from 319 PCR replicates for beluga and 355 PCR replicates for Eulachon, coho, and chinook after discarding samples with a coefficient of variation greater than 1.5 across technical replicates. We considered a detection within a bottle any Ct less than 40 cycles across technical replicates, regardless of coefficient of variation. For the coho assay, we observed 213 detections with a mean concentration of 31,070 ± 47,878 copies/L (max=338,892 copies/L) across detections (Table S3). For the Chinook assay, we observed 106 detections with a mean concentration of 2,068 ± 2913 copies/L (max=15,530 copies/L) across detections (Table S4). For the eulachon assay, we observed 204 detections with a mean concentration of 17,464 ± 72,867 copies/L (max=627,824 copies/L) across detections (Table S5). For the beluga assay, we observed 56 detections with a mean concentration of 3,140 ± 12,012 copies/L (max=51,775 copies/L) across detections (Table S6). Due to uniformly low estimated DNA concentrations for beluga whales and high limits of quantification (Appendix 3), we summarized beluga whale observations in terms of detections.

### Detecting Cook Inlet beluga whales with eDNA

#### Twentymile River

We detected beluga whales in eDNA samples using both species-specific qPCR and MiFish metabarcoding assays (Figure 3). For qPCR results for Cook Inlet beluga whales, we only analyzed presence absence occurrence data and did not use results from the above model, as overall performance was poor with most positive amplifications near the limit of detection of 1 copy per reaction (Appendix 3). We found both qPCR and metabarcoding positively detected beluga in 70% of samples while 23.4% of samples were only detected by qPCR only and 6.2% only by metabarcoding only (Figure 4).

**Figure 3.**
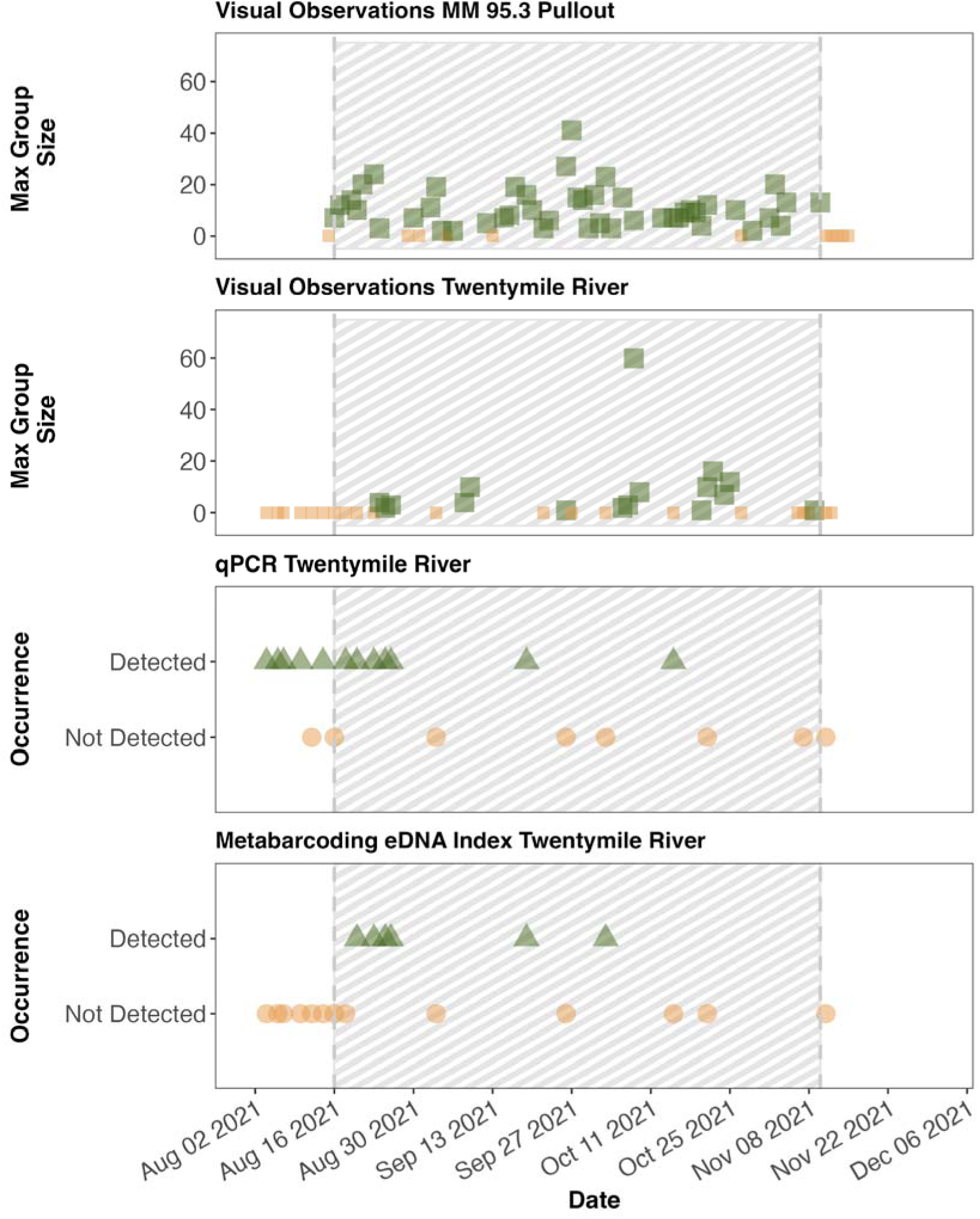
Visual and genetic detection of beluga whales in the vicinity of Twentymile River, upper Cook Inlet, Alaska. Detections are green triangles and absences are brown circles. The hashed area indicates the period of beluga whale occupancy based on visual detections. Both species specific beluga whale qPCR and MiFish *12S* metabarcoding assays detected belugas in the vicinity of Twentymile River on multiple occasions over the three month sampling period. Beluga whales were detected using molecular genetic assays prior to the first visual detection of beluga whales in the vicinity of Twentymile River in August 2021.

**Figure 4.**
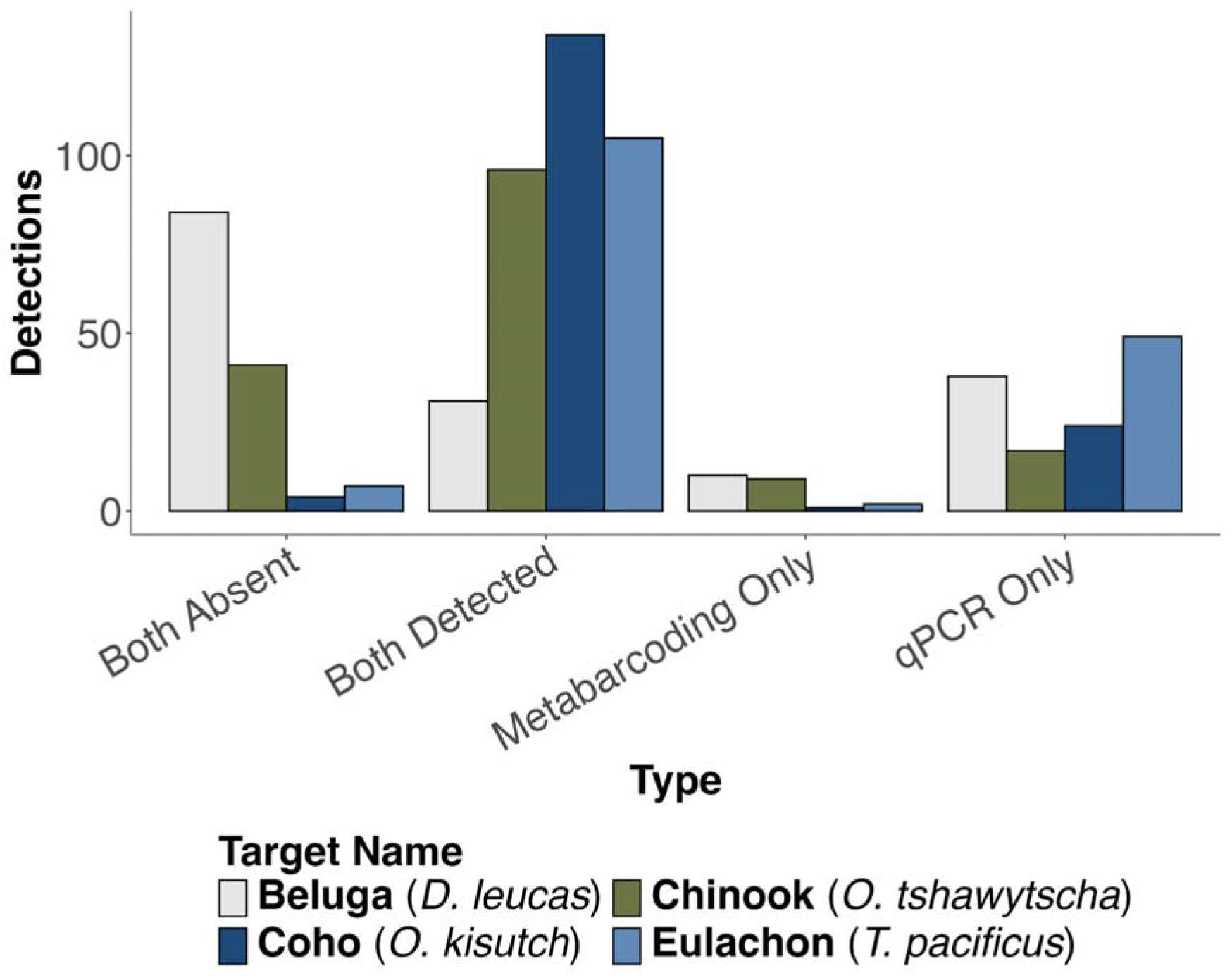
Comparison of qPCR and metabarcoding detections for beluga whales, Chinook (*O. tshawytscha)*, coho (*O. gorbuscha*), and eulachon (*T. pacificus*) across all observations. Beluga whales (*D. leucas*) were more frequently detected by qPCR than metabarcoding. In contrast, all three fish species were most frequently detected with both qPCR and metabarcoding.

During the study period, beluga whales were first visually observed in Turnagain Arm at Bird Rock on August 5th (S. Steinhert *pers. comms*) prior to the start of AKBMP visual observations at Mile Marker 95.3. The visual AKMBP observations started on August 15th, 2021 with the first visual occurrence on August 16th, 2021 at Mile Marker 95.3. Cook Inlet Belugas were first observed actively swimming up Twentymile River on August 21st, 2021. Beluga whales were last observed in the vicinity of Twentymile River on November 9th, 2021 and at Mile Marker 95.3 November 10th, 2021. We used these positive visual detections to define the period of beluga occupancy. During the defined period of occupancy, beluga whales were detected in 69.2% of visual monitoring survey time points at Twentymile River. During the same time period, beluga whale DNA was detected in 46.2% of eDNA samples using metabarcoding and 64.3% of eDNA samples using qPCR (Figure 3).

Both eDNA methods detected beluga whales in the vicinity of Twentymile River before whales were visually observed at either Mile Marker 95.3 or Twentymile River monitoring stations (qPCR on August 4th, and metabarcoding on August 20th). In particular, qPCR approaches detected Cook Inlet beluga whales prior to the first visual detections at the 95.3 Mile Marker (18 km from the eDNA sampling location) on August 16th. Multiple AKBMP volunteers confirmed that no beluga whales were visually observed inshore of the 95.3 Mile Marker during August 4th and August 14th, during *ad hoc* monitoring sessions taken prior to the official start of AKBMP monitoring (S. Steinher *pets. comms*). These results suggest the potential transport of beluga whale DNA by the incoming tidal bore. In contrast, metabarcoding approaches only detected beluga whale during the period in which beluga whales were observed in the vicinity of Twentymile River, albeit more inconsistently than visual surveys overall.

#### Kenai River

In spring 2021, AKBMP monitoring at Kenai began on March 15, the first visual sighting occurred on March 25, the last visual sighting occurred on April 30, and monitoring continued until May 29. In fall 2021, monitoring began on August 15, the first visual sighting occurred on September 3, and the last visual sighting occurred on November 15, which was also the final monitoring session. Cook Inlet beluga whales were predominantly observed in Kenai River from visual surveys in fall (September 3rd, 2021 to November 15th, 2021, detection rate of 67.2% per observation session) and spring (March 25th, 2021 to April 30th, 2021, detection rate of 70.3% per observation session), but were absent with zero visual detections during summer visual surveys (May through September; Figure 5). eDNA also detected beluga whales in fall and spring, with qPCR detecting beluga whale in 100% of sampling events in fall and spring while metabarcoding detected beluga whale in 66.7% of sampling events in fall and 100% in spring. eDNA also frequently detected beluga whales outside of the visually defined period of occupancy in all but two of 14 months sampled (June and August). Similarly, eDNA metabarcoding detections were positive for beluga whales in 11 out of 14 months with only July, August, and October having no detections. We note that in late spring and summer months where visual surveys did not detect beluga whales only one or two technical or bottle replicates detected beluga whales, suggesting that the occurrence of beluga whales in Kenai River during this time period was very rare.

**Figure 5.**
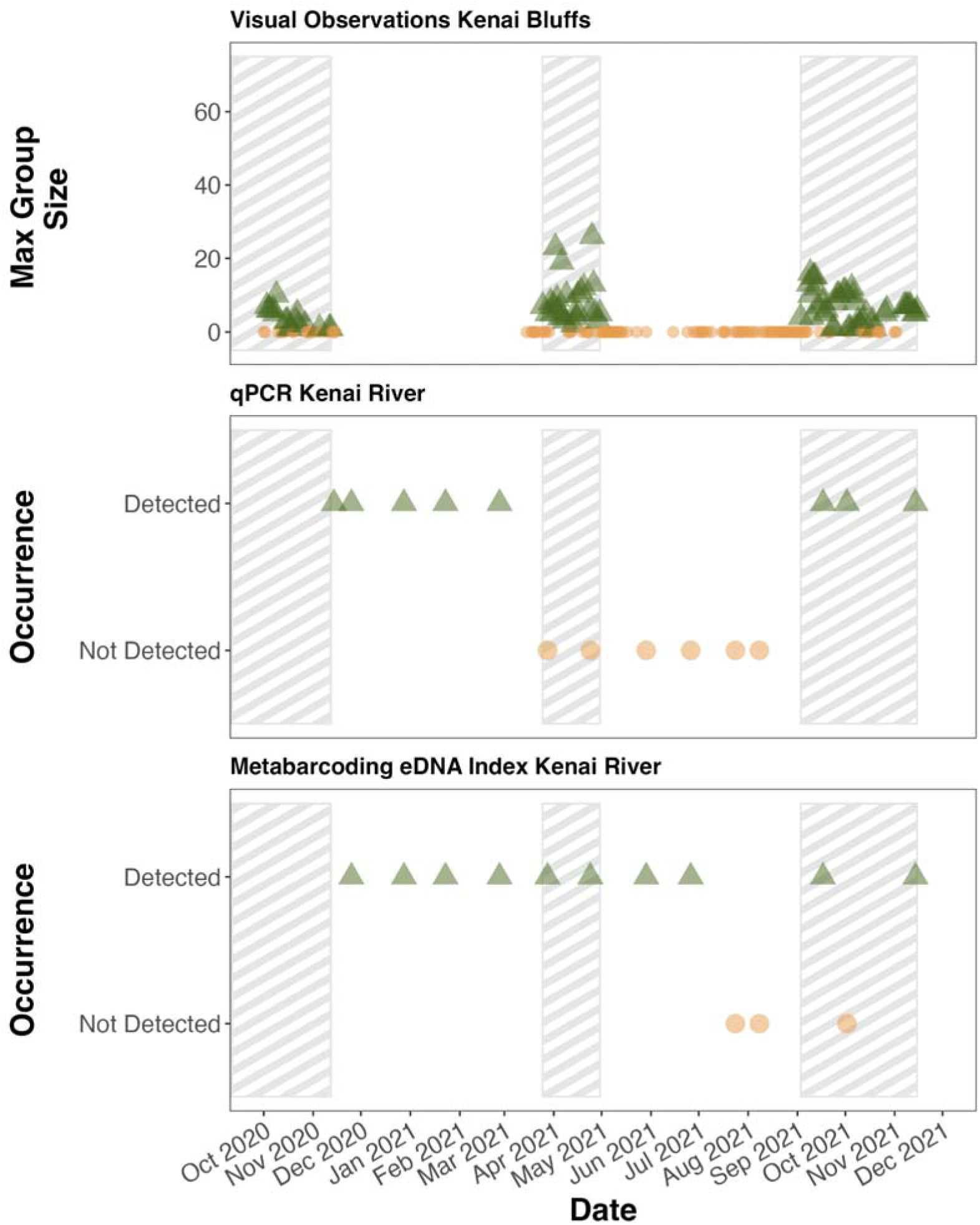
Cook Inlet beluga whale detections in Kenai River, Alaska. Detections from both the dock and bridge site are green triangles and absences are brown circles. Both species-specific beluga whale qPCR assays and MiFish *12S* metabarcoding detected beluga whales in the Kenai River system.

### Quantitative Metabarcoding Reveals Co-Occurrence of Beluga Whales and Salmon Prey in the vicinity of Twentymile River, AK

We focused our analyses on temporal changes in salmonid abundance given that salmonids are key prey source for Cook Inlet beluga whales and comprised a strong majority (83.1%) of total observed DNA reads (Quakenbush et al. 2015). In the vicinity of Twentymile River, DNA copies per L of coho salmon (*O. kisutch*) more than doubled during beluga whale occupancy from the period prior to beluga whale occupancy (Figure 6). In contrast, we found beluga whale migration into the river corresponded with a 85.6% decrease in the abundance of Chinook salmon DNA concentration (*O. tschwaytscha*). The concentration of pink (*O. gorbuscha*), chum (*O. keta*), steelhead (*O. mykiss*), or sockeye (*O. nerka*) DNA were similar before and during the period of beluga whale occupancy in the same region. Pink salmon were the most abundant species across the entire sampling period in the vicinity of Twentymile River with the highest variability across and within sampling time points and exhibited a marked decline throughout the period of beluga whale occupancy. In contrast, coho abundance increased and largely remained abundant during the rest of the sampling period (Appendix 6).

**Figure 6.**
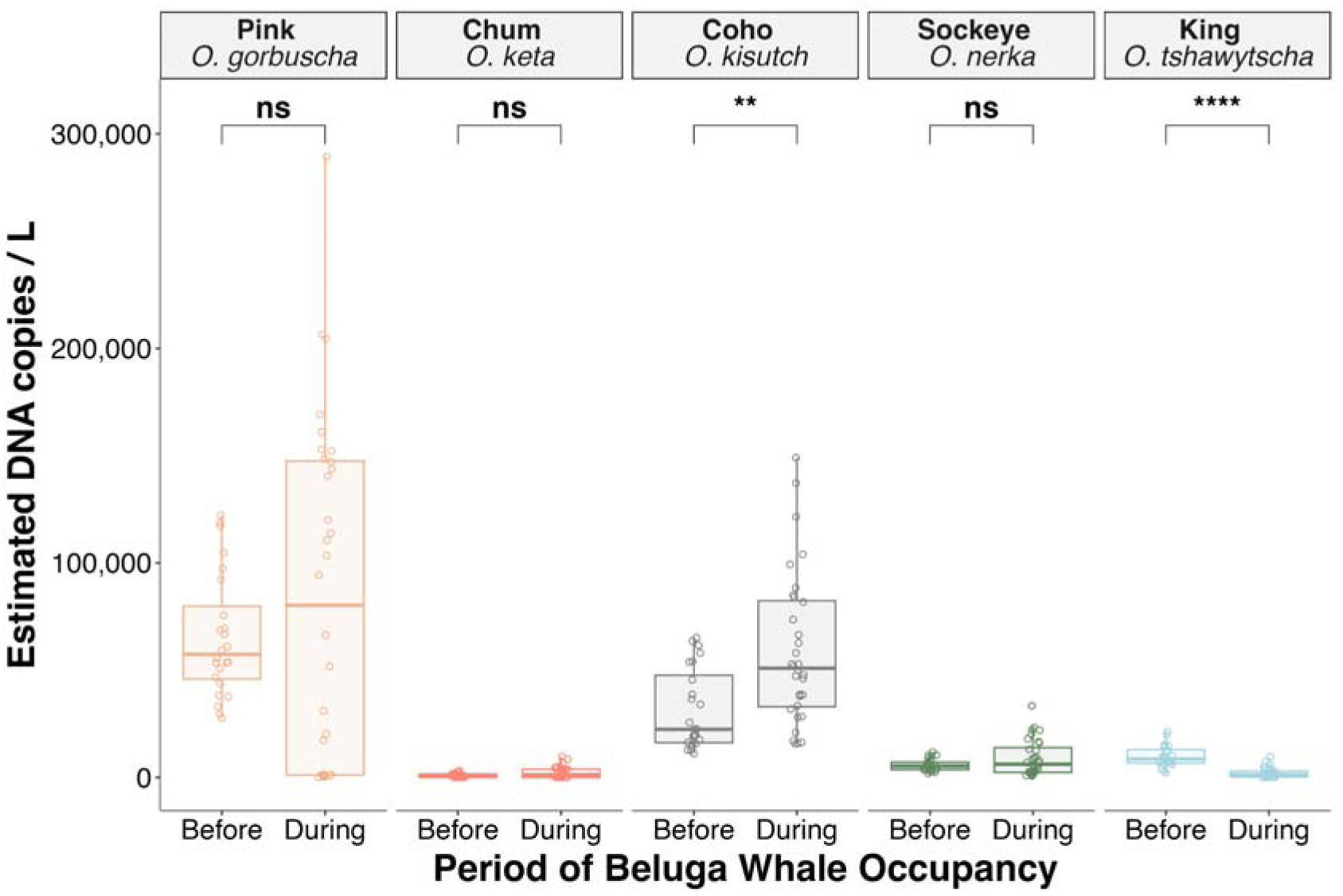
Salmon abundance in the vicinity of Twentymile River as measured by DNA copies per liter of seawater in surface eDNA samples. Median coho salmon (*O. kisutch*) concentrations were 228% higher when beluga whales occupied Turnagain Arm, than before beluga whales were observed in the estuary. The highest DNA concentrations were from Pink salmon (*O. gorbuscha*) throughout the study period and showed a clear peak and decline prior to when beluga whales were observed in the estuary (Appendix 6) with no significant difference in abundances across time periods (p>0.05). Boxplots represent median, interquartile range, and whiskers extend to maximum and minimum values. Significance values derived from pairwise Kruskal-Wallis comparisons of estimated DNA abundance are shown (ns: p > 0.05, *: p <= 0.05, **: p <= 0.01, ***: p <= 0.001, ****: p <= 0.0001).

### Evidence of Swamping in Metabarcoding Results

If non-detections of metabarcoding data were occurring due to an overwhelming signal to noise ratio, we would expect to observe non-detections in metabarcoding data from samples with high concentrations of DNA as measured by qPCR. Here we refer to these non-detections as “swamping” of a metabarcoding signature in which a comparatively weaker signal is overwhelmed by a more dominant signal of DNA. We observed 137 instances in which fish species were detected in qPCR data, in the absence of metabarcoding reads (Figure 7). These metabarcoding non-detections occurred for coho, Chinook, and eulachon in Kenai River and Chinook and eulachon on Twentymile River. Here we argue that swamping arises due to the competitive PCR dynamics and compositional constraints of metabarcoding sequencing, resulting in over amplification bias of dominant taxa and unexpected non-detections of comparatively rare taxa in mixed-community metabarcoding. Because metabarcoding generates strictly compositional data, the exponential over-amplification of such dominant taxa can result in a complete dropout of less competitive sequences, leading to false negative detections.

**Figure 7.**
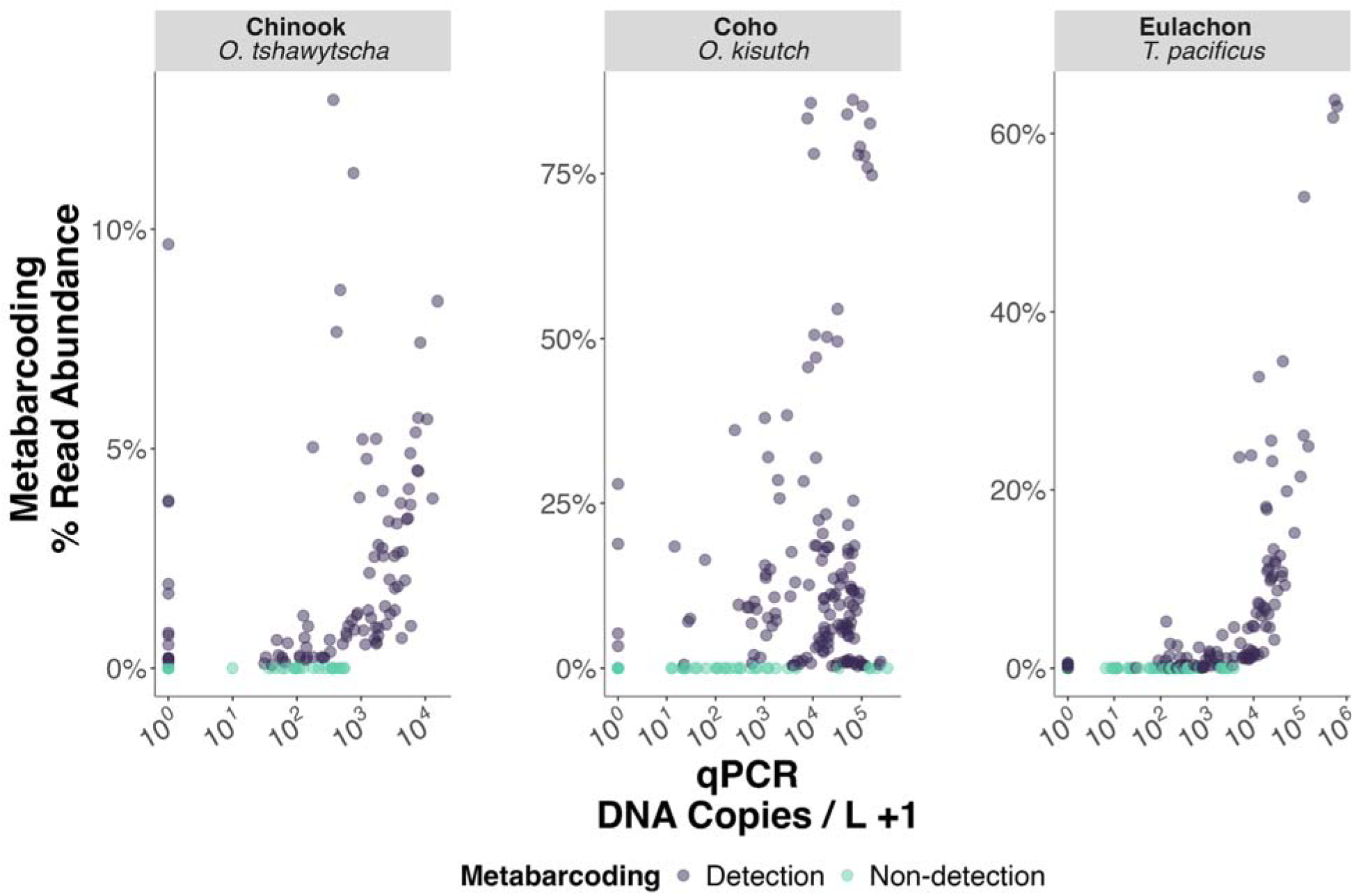
Evidence of swamping of metabarcoding signatures. We observed non-detections in metabarcoding data (green) when qPCR results from the same samples measured tens of thousands of copies per liter. Each point represents qPCR derived DNA concentrations on the x axis and metabarcoding derived percent read abundance for a given species in a given sample. Notably, a sample with 338,892 coho copies per L resulted in a non-detection in metabarcoding. A non-detection is defined as zero observed metabarcoding reads assigned to that species across all bottle and PCR replicates.

Metabarcoding non-detections for coho occurred in a total of 35 instances. The most notable instances of swamping (n=7) occurred in June and July 2021 where sockeye salmon were on average 65% of all reads (max 89.4%, min 16.6%) of MiFish metabarcoding reads in each sample. qPCR concentrations successfully detected coho in these samples, ranging from 3,063-338,892 copies/L (Table S3).

### Spatial and Temporal Variation of eDNA Signatures in the Kenai River

Variability in fish community species composition and dispersion in the Kenai River was strongly driven by sampling date, and to a lesser degree, site (Figure 7, PERMANOVA, p =0.001). Temporal variability across months explained the greatest variance in fish assemblages (52.8%) compared to site (3.3%), depth (0.5%), and biological replication (e.g., distinct sample bottles) (0.4%, not significant). The interaction of sampling date with site (5.6%), depth (6.0%), and biological replication (4.5%) explained additional variance in captured eDNA signatures. We also found a significant correlation between time and the homogeneity of dispersion of captured eDNA signatures across biological replicates, indicating that the variance among eDNA samples itself was correlated with seasonal changes. Here we saw eDNA communities collected in August were the most similar while eDNA samples in April were the most dispersed (Appendix 7). We observed no difference in dispersions between sites on Kenai River (Homogeneity of Dispersons, p> 0.05).

Ordination of Kenai River eDNA samples shows an annual pattern of fish community assemblages oriented like a seasonal clock with samples taken in overlapping seasons looking more similar to each other (Figure 8). These results align well with the known cycle of anadromous fish migrations that result in periodic occurrences of species assemblages within the Kenai River throughout the year (<u>Weiner, 1998</u>). Together these results strongly suggest that sampling date was the strongest predictor of fish community assemblage patterns in this anadromous fish dominated riverine fish community.

**Figure 8.**
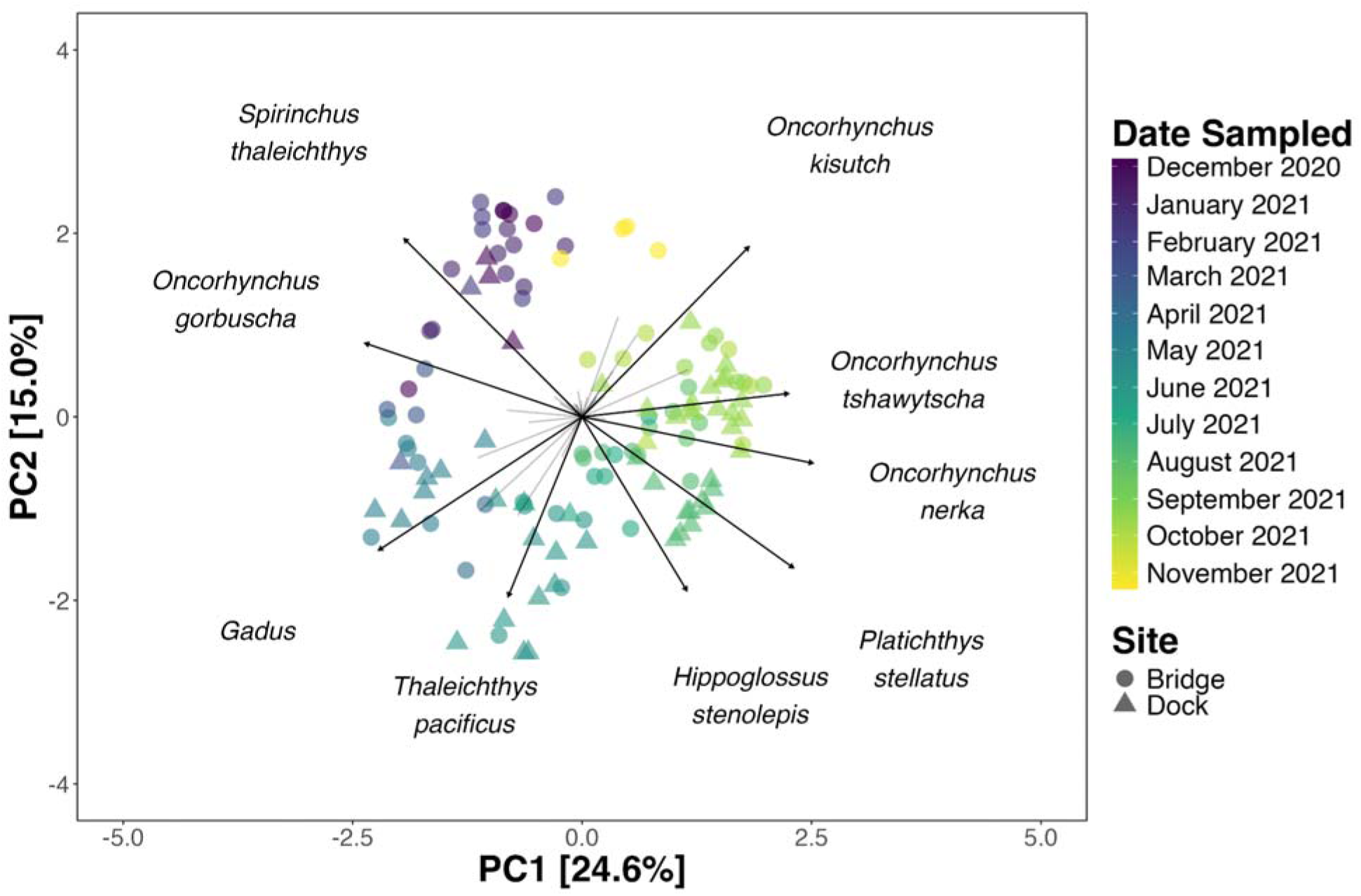
eDNA signatures vary by sampling date. We see clear seasonal annual patterns of fish communities within the Kenai River. Principle coordinates analysis ordination plot of Aitchison dissimilarities of eDNA reads.

From our qPCR results, we observed the highest qPCR derived absolute abundances of eulachon (*Thaleichthys pacificus*) at the end of May (297 ± 709 copies/L), after the visually defined period of beluga whale occupancy in the spring. Cook Inlet beluga whale were visually observed in March and April when eulachon abundance was two orders of magnitude lower (March: 0.82 ± 0.94 copies/L; April: 2.1 ± 2.3 copies/L; Appendix 5). During March and April we observed higher proportional abundance of gadids, *Cottus* sp., *Sebastolobus* sp., chum, and sablefish (*Anoplopoma fimbria*). These results suggest that beluga whales were occupying Kenai River outside of the period of eulachon abundance.

## Discussion

Interrogating eDNA samples with both fish metabarcoding and qPCR assays revealed a dynamic prey community in Cook Inlet. Changes in beluga whale occurrence and concurrent changes in the composition of fish communities in both Twentymile and Kenai estuaries highlight the utility of eDNA samples for characterizing changes in habitat use by cetaceans and their putative prey. Leveraging both visual survey methods and near-shore eDNA samples, we demonstrate the value of quantitative metabarcoding eDNA approaches to provide critical insights into the co-occurrence of Cook Inlet beluga whales and temporal changes in the abundance of key fish prey species. Together, these results lay the foundation for future research efforts to characterize Cook Inlet beluga whale spatial and temporal foraging patterns needed to inform the management and recovery efforts of this critically endangered species.

### Detecting Cook Inlet Beluga Whales with eDNA

We demonstrate the success of a species-specific beluga whale qPCR assay and MiFish *12S* metabarcoding for detecting Cook Inlet beluga whale (Figures 3 & 4). Previous results have found that eDNA approaches often have a higher false negative rate for the detection of marine mammal species (Pinfield et al., 2019; Suarez-Bregua et al., 2022). In contrast, here we found that our qPCR assay consistently detected Cook Inlet beluga whale largely on par with visual surveys. Interestingly, despite a lower rate of detection, metabarcoding results better matched visual observations on Twentymile River as opposed to qPCR approaches which frequently detected beluga whales when they were only observed over 18 km away from Twentymile River suggesting the transport of shed DNA molecules over significant distances in this dynamic, high flow environment. Together, these results suggest that species-specific qPCR approaches are more sensitive than MiFish *12S* metabarcoding approaches. This is likely due to a combination of both lower amplification efficiency and rarer abundance of marine mammal DNA relative to teleost fish (Shaffer et al., 2025) aligning well with proposed mechanistic understanding of the compositional nature of metabarcoding (Gloor et al., 2017; McLaren et al., 2019; Shelton et al., 2023; Silverman et al., 2021). Our results provide further illustration of how qPCR and metabarcoding approaches encompass distinct views of eDNA signatures in the marine environment. Given that observed metabarcoding detections are dependent on both the true abundance of Cook Inlet beluga whale eDNA concentrations and the relative abundance compared in a given sample, it makes sense that Cook Inlet beluga whales were only observed by metabarcoding approaches when they were physically observed in the vicinity of Twentymile River and were likely contributing a greater relative proportion of total DNA in the water column. In contrast, qPCR detections are independent from the sampled DNA of other species in the water column and thus was able to detect proportionally rare molecules of Cook Inlet beluga whale DNA. In addition, qPCR hydrolysis probe assays (i.e. Taqman assays) are generally more sensitive than metabarcoding approaches (Wood et al., 2019) and in this case, target smaller DNA fragments, both increasing the potential for positive detections in more degraded eDNA samples.

### Transport of eDNA in Highly Dynamic Tidal Environments

We provide evidence for the transport of eDNA over ten kilometers during a tidal bore event in an estuary system (Figure 1). This represents one of the largest (by distance) measures of eDNA transport in a marine environment in the literature to date, on par with observations of eDNA transport in high transport aquatic and marine environments (Baetscher et al., 2024; Deiner & Altermatt, 2014; Jo & Yamanaka, 2022b; Kutti et al., 2020; D. Shea et al., 2022; Xiong et al., 2025). Both Twentymile and Kenai Rivers experience exceptionally large tidal cycles driving dramatic water mixing, facilitating the transport of eDNA on the scales of tens of kilometers (Fissel & Lin, 2022; Oey et al., 2007; Wang & Yang, 2020). Given the low likelihood of false negative visual detections and repeated non-detections from visual and eDNA surveys prior to whales being observed in Turnagain Arm, detections due to resuspension of beluga whale DNA from river sediments seems unlikely. Furthermore, such observations of dramatic marine eDNA transport were observed on ten separate sampling events. We detected beluga whales five times between August 3rd and August 16th in the vicinity of Twentymile River prior to the start of the visual surveys, but while *ad hoc* observations did not observe belugas inshore of Bird Point (SSteinhert *pers. comms*). In some instances whales may have travelled closer to Twentymile River than Mile Marker 95.3. However, no whales during this period were ever observed within 4 km of the mouth of Twentymile River (the limit of visual observation), suggesting DNA transport of at least this distance. Regardless of precise transport distance, our results provide strong evidence for transport of eDNA signatures on the scale of tens of kilometers in a highly dynamic tidally driven system.

In addition to physical transport from the tidal bore in Turnagain Arm, both Cook Inlet beluga whale behavior and cold temperatures likely aided in the facilitation of eDNA transport observed here (<u>Hashizume, 2015</u>). Previous observations have found correlations between tidal height and the occurrence of Cook Inlet beluga whales in Turnagain Arm, with an increased frequency of whale observations during tides greater than 10 m that are associated with large tidal bores (Ezer et al., 2008; Moore et al., 2000). Such “surfing” behavior immediately swimming along the tidal bore likely helped facilitate transport of DNA molecules and may suggest that beluga whale eDNA dynamics may be distinct from other species in this system that do not display such behaviors. In addition, previous work has demonstrated eDNA degradation to be slower in colder waters (Joseph et al., 2022; McCartin et al., 2022; Strickler et al., 2015).

Our observations support the hypothesis that the ecological integration of eDNA signatures is, in part, a function of the hydrodynamics of a given system (Andruszkiewicz et al., 2019; Harrison et al., 2019; Shogren et al., 2017; Xiong et al., 2025). From this hypothesis, we expect that eDNA transport is greater in areas with increased advection and flow rates. This mechanistic hypothesis helps explain both observations of multi-kilometer scale transport observed here and elsewhere (Kutti et al., 2020), as well as from previous studies that have found strong differences in eDNA signatures on the scale of tens of meters in comparatively calm, low transport nearshore environments (Monuki et al., 2021; Port et al., 2016; M. M. Shea & Boehm, 2024). These results highlight that exceptional eDNA transport can occur in highly dynamic marine environments. These findings echo the need for careful consideration of localized physical oceanography in the interpretation of eDNA signatures.

Observations of spatial, temporal, and depth variation here provide important information for marine eDNA sampling design. Given the scale of eDNA transport on the order of multiple kilometers within Cook Inlet, we expect eDNA samples collected near the mouth of Twentymile River to represent a spatially integrated signature of Turnagain Arm. Our results indicate that it is unlikely that eDNA samples would be capable of capturing differences in fish assemblages at fine spatial scales within a river system.

Our results suggest that conducting eDNA sampling on a few key regions within Cook Inlet (e.g., Kenai River, Turnagain Arm, Chickaloon Bay, Susitna River, Beluga River, and Knik Arm) with frequent sub-weekly to weekly sampling would provide the spatial and temporal resolution needed to link beluga whale and prey co-occurrence and provide critical information on putative prey fields throughout their critical habitat. Moreover, the coarse spatial integration of eDNA signatures in Cook Inlet has implications for a broad range of potential eDNA applications. For example, we would not expect to be able to evaluate differences in fish communities between sites with offshore energy production or port modifications within Cook Inlet that may be expected to have localized impacts on the scale of meters to a kilometer. However, the observed resolution of eDNA signatures is ideal for capturing regional and temporal differences in putative prey abundance for Cook Inlet beluga whales across 10km wide estuary deltas that are situated between 40 and 500 km apart from each other (McHuron et al., 2023).

### Improving Quantitative Metabarcoding Estimation Approaches

Our results provide generalizable insights informing effective quantitative metabarcoding eDNA applications that build off of integrating qPCR and metabarcoding observations. We provide evidence for the importance of identifying an effective qPCR target to anchor absolute abundance estimates across eDNA samples. This can be challenging in practice when species communities fluctuate across samples. The variance in qPCR observations increases for low concentrations (Karlen et al., 2007; McCall et al., 2014; Ruijter et al., 2009; Shelton et al., 2019, 2023) and thus accuracy of estimates of total DNA decline when an anchor species is rare. For example, coho, eulachon, and Chinook abundances were low or absent in winter months on the Kenai River, making estimates of the total DNA within a community challenging. The solution to this problem is to ensure that appropriate anchor species are chosen for each sample. In Cook Inlet, such an approach would require the utilization of multiple qPCR assays for systems with dynamic fluctuations in abundant species with ∼8 orders of magnitude changes in fish biomass (kg) across seasons (Alaska Department of Fish and Game, n.d.; Gates et al., 2024). Here we recommend that future quantitative metabarcoding approaches in Cook Inlet riverine systems target sockeye, pink, and chum salmon in addition to Chinook and coho implemented here to ensure that all abundant species are captured. The need to include multiple qPCR targets to anchor quantitative metabarcoding approaches extends to other systems with large variation in abundance. In such instances, a tractable solution would be to conduct metabarcoding first and use the generated data to select anchor species across the full data set.

In addition to having accurate absolute abundance estimates, our work emphasizes that of others highlighting the need for comprehensive mock communities to apply the proportional bias correction to metabarcoding data. Here, we only successfully derived PCR amplification efficiency for five salmonid species due to a mismatch in generated mock communities and observed species. Without estimates of amplification efficiency, we can not accurately estimate true underlying relative abundance between taxa and therefore unable to accurately estimate absolute DNA concentrations for other taxa.

Fortunately, developing mock communities for freshwater and marine fishes in Cook Inlet and its tributaries is relatively straightforward for such a low diversity environment (<100 species). Although developing mock communities in other systems may be challenging, a single dedicated sequencing run of mock communities could easily incorporate all known fish prey targets and their congeners. We strongly recommend future quantitative eDNA metabarcoding approaches carefully consider their target species and existing ecological knowledge to make informed decisions in generating mock communities. Developing mock communities for hyperdiverse environments remains a challenge for the field and will likely require alternative mechanisms such as variable PCR cycling or unique molecular identifiers (see Shelton et al., 2023) to enable accurate estimation of amplification efficiencies needed to derive quantitative metabarcoding estimates.

This work highlights the value of combining metabarcoding and qPCR approaches to implement more robust quantitative eDNA approaches across multiple species, expanding the scientific capabilities and robustness of ecological interpretation. Here we accounted for amplification bias in our metabarcoding data to enable absolute abundance calculations for multiple species (Allan et al., 2023; Shelton et al., 2023).We focused our analyses on salmonids which made up the vast majority of observed DNA sequencing reads and were included in our mock communities.

We note that this model did not account for all sources of variation that may affect recovered eDNA concentrations. Previous work has demonstrated that total DNA concentration in a given liter of water is expected to be a function of the total underlying abundance of a species in the environment, the shedding rates, degradation rates, fate and transport of DNA, and laboratory biases, namely PCR amplification (Harrison et al., 2019; Shelton et al., 2019, 2023). Shedding and degradation rates have been demonstrated to be a function of species specific physiological and behavioral traits and microbial activity in the environment (Andruszkiewicz Allan et al., 2021; Barnes et al., 2014; Beattie et al., 2023; Joseph et al., 2022; Thalinger et al., 2021; Waters et al., 2024; Zhao et al., 2023). Given the relative similarity in physiology of species within the same genus, sampled concurrently in the same environment, we expect shedding and degradation rates to be largely similar across the salmonids analyzed here. Thus we argue that our analyses of eDNA concentrations across species is largely reflective of the changes in underlying abundances. These results are supported by other applications of metabarcoding and qPCR observations of salmon abundance (Benoit et al., 2023; Levi et al., 2019; Robinson et al., 2023; Shelton et al., 2019; Tillotson et al., 2018).

### Evidence of Swamping of eDNA Metabarcoding Signals

Here we provide evidence that competition in the metabarcoding PCR reaction between DNA molecules of multiple species can result in swamping. An eDNA sample collected on the Kenai River in July near the peak of the sockeye run was comprised of 84% sockeye salmon reads and zero coho reads, despite having a qPCR estimated absolute coho concentration of 300,000 copies/L. Amazingly, the metabarcoding results only detected one other species, Dolly Varden (*Salvelinus malma*) at 16% of total reads, and failed to detect coho entirely, despite this eDNA sample having the highest yield of coho DNA concentration observed in this study. This observed swamping of eDNA signatures aligns with previously developed quantitative metabarcoding and signal-to-noise frameworks (Allan et al., 2023; Gleason et al., 2023; Gloor et al., 2017; Gold et al., 2023; McLaren et al., 2019; Shelton et al., 2023; Silverman et al., 2021; Wood et al., 2019)). Our results demonstrate that the lack of a metabarcoding detection of species does not preclude a high absolute abundance of the species when other more abundant species are cooccur (Gloor et al., 2017) that the application of eDNA metabarcoding approaches for characterizing non-dominant species will be challenging to interpret in high biomass environments (e.g., salmon runs, large schools of fish, broadcast spawning events) for both quantitative estimates and species richness.

Ultimately, the occurrence of swamping in metabarcoding data highlights the need for careful ecological interpretation of compositional amplicon sequencing signatures alone (Deiner et al., 2017; Shelton et al., 2022). These results reinforce the value of pairing amplicon sequencing approaches alongside other absolute observational data sources (qPCR, visual observations, etc.) to interpret compositional amplicon sequencing (Taberlet et al., 2018).

### Beluga whale occurrence in the vicinity of Twentymile River is Correlated with Coho Salmon Run Timing

Cook Inlet beluga whales occurred in the vicinity of Twentymile River when coho salmon abundances were highest. These results align with previous observations of other beluga whale populations which seasonally utilize freshwater habitats to target anadromous fish migrations (Breton-Honeyman et al., 2016; Castellote et al., 2020, 2021; Nelson et al., 2018; Quakenbush et al., 2015; Watt et al., 2016) align with traditional ecological knowledge shared by local residents fishing on Twentymile River in 2021 (Paul Crews *pers. comm.*).

It is important to recognize that such a co-occurrence framework between predator and putative prey species can be fraught with challenges. For example, presence-absence data lacks sufficient information for ecological inference of interactions (Blanchet et al., 2020) as species can co-occur in space for a myriad of reasons, but particularly environmental filtering in which habitat constraints are driving co-occurrence patterns (e.g., have similar thermal niches) (Dormann et al., 2018). Positive co-occurrences are more reasonably interpreted as a predator-prey relationship when observations are made in the context of active foraging, in a habitat that is only used by a predator when it is exhibiting foraging behavior, and predator occurrence is being compared to changes in abundance of prey targets (Amir et al., 2022; Carroll et al., 2019; Ford & Goheen, 2015). In such a context, it is clear that foraging behavior, not environmental filtering, is driving such positive co-occurrence patterns.

Here, we argue that co-occurrence patterns of beluga whales and putative salmon prey in the vicinity of Twentymile River satisfy these criteria and therefore changes in eDNA based abundance of coho can be reasonably interpreted as predator-prey associations. During this study we observed behavior indicative of beluga whale foraging in the vicinity of Twentymile River, repeatedly associated with high tides across a three month span. Foraging behavior is typically characterized by repeated, short (<10 minutes), and shallow (<10m) water dives with slow transit time, consistent with observed behavior on Twentymile River (Castellote et al., 2020; Goetz et al., 2012; McGuire, Himes Boor, et al., 2020). In addition, previous work in Cook Inlet has found a strong positive seasonal association with Cook Inlet beluga whale occurrence, feeding calls, and salmon species on the Susitna and Chickaloon rivers, providing strong evidence for active foraging of salmon species in similar riverine habitats within Cook Inlet (Castellote et al., 2020, 2021); beluga whales are not known to readily utilize enclosed riverine habitats unless actively foraging, as shallow water habitats increase the risk of stranding. Furthermore, Cook Inlet beluga whales only intermittently utilize this habitat both seasonally (spring and fall) and are more frequently observed entering Twentymile River on tides greater than 10 m, when there is a more substantial connection of water between the mouth of the river and Turnagain Arm (McHuron et al., 2023; Moore et al., 2000). Lastly, over a decade of visual observations of Cook Inlet beluga whales have documented active pursuit of coho salmon (McGuire, Himes Boor, et al., 2020) and they have been positively identified as prey items in Cook Inlet beluga whale stomachs (Quakenbush et al., 2015).

### Beluga Whale Co-Occurrence Observations on the Kenai River

Our low sampling frequency and incomplete mock communities prevented the accurate quantitative characterization of broad fish community abundances in Kenai River. However, qPCR results provide quantitative data for Chinook, coho, and eulachon quantitative co-occurrences. In addition, our non-bias corrected eDNA metabarcoding provides data for generating hypotheses of potential putative prey to be examined in future studies.

In the spring of 2021, AKBMP volunteers visually observed Cook Inlet beluga whales in April and May in the Kenai River. Here we found no evidence of co-occurrence of beluga whale and Chinook, coho, nor eulachon during this time period. All three species increased in abundance at the end of May and June after beluga whales were observed in the Kenai River. Importantly, the mismatch between the observed maximum eulachon concentration and beluga whale occurrence are particularly notable as beluga whale have been previously documented to forage on such large annual migrations of eulachon and other osmerid smelt species in Twentymile and Susitna rivers. However, with only monthly sampling our ability to resolve such a mismatch is limited. Higher temporal sampling is needed to better understand spring beluga whale foraging patterns in Kenai River. How limited results hint at the possibility that beluga whales are likely not targeting spring eulachon runs as a prey source. Instead, during April and May we detected *Gadus* sp. and Sablefish (*Anoplopoma fimbria*) from *MiFish 12S* metabarcoding (Appendix 6). Both species use estuaries and shallow marine environments during juvenile life stages and have also been identified as prey targets for beluga whales in other regions (Abookire et al., 2000). Future work is needed to determine beluga whale foraging patterns and targets in the Kenai River during spring months.

Monthly Kenai River eDNA data also detected beluga whales in winter months alongside metabarcoding detections of longfin smelt (*Spirinchus thaleichthys),* sculpin (*Cottus* sp.), three spined stickleback (*Gasterosteus aculeatus*), Pacific sandlance (*Ammodytes personatus*), coho (*O. kitsuch)*, and pink salmon (*O. gorbuscha*). These winter detections of Cook Inlet beluga whale eDNA in the Kenai River suggest either beluga whales may utilize the river even when it is ice covered in winter months (Dutton et al., 2012; Ezer et al., 2008; Moore et al., 2000) or alternatively represent tidal transport of eDNA from the river mouth into the river (<u>Kumar et al., 2024</u>). Such observations point to the need to increase winter eDNA sampling efforts when Cook Inlet beluga whales may be swimming underneath the frozen river surface, leveraging better-designed quantitative metabarcoding approaches than applied here.

Lastly, in the last few decades Cook Inlet beluga whales have been largely absent from the Kenai River during the summer and do not return until August (AKBMP unpublished data, Kumar et al 2024). However, eDNA observations did detect beluga whale during summer months, albeit inconsistently across technical PCR and biological sample replicates. This may indicate either there were few individuals that did occur, or more likely, eDNA advected from the mouth of the estuary or nearby coastal habitat into the river. Historical observations have noted that beluga whale frequented Kenai River in the summer in the same time period (Dutton et al., 2012; Huntington, 2002). Chinook and sockeye runs historically peak during these months which are known prey sources. However, anthropogenic activity is also at its highest during this period, with substantial recreational fishing and commercial vessel traffic (Brewer et al., 2023). Future surveys pairing acoustic observations of belugas and anthropogenic noise alongside eDNA during spring, summer, and fall months would provide much needed data to help understand the interplay between beluga occurrences, putative prey fields, and anthropogenic noise on Kenai River (Kumar et al., 2024).

### Informing Future Prey Survey Efforts

A major challenge of marine mammal conservation and management is the development of robust, paired data streams on population dynamics, marine mammal movements, and trophic ecology (Avila et al., 2018). Such information is needed to better understand the relative impacts of multiple stressors on coastal marine mammal species and support informed conservation and management efforts (Booth et al., 2020; Nelms et al., 2021). For Cook Inlet beluga whales, existing aerial, acoustic, tagging, and visual surveys have provided valuable information on their seasonal movements and the landscape of anthropogenic noise (Castellote et al., 2021; Goetz et al., 2023). This extensive body of research has shown that beluga whales display seasonal movements across Cook Inlet associated with optimal seasonal foraging of anadromous fish migrations (Castellote et al., 2020, 2021; McHuron et al., 2023). However, concurrent characterization of Cook Inlet beluga whale foraging ecology is limited because of a lack of empirical data on prey consumption and prey fields, particularly in smaller, understudied rivers. By developing concurrent observations of beluga whale occurrences, anthropogenic disturbance, and their prey landscape across their habitat, we can understand if such disturbances are negatively impacting their ability to utilize prey resources (McHuron et al., 2023). Our results provide the groundwork for leveraging quantitative eDNA approaches for future studies of Cook Inlet beluga whale foraging patterns to provide robust data on year-round prey availability in Cook Inlet.

Our work strongly suggests that the application of a basin wide eDNA monitoring network at appropriate spatial scales could provide important insights into the seasonal foraging patterns of Cook Inlet beluga whales across the extent of their range. Here we advocate that incorporating a few minor additional methodology changes along with a more robust weekly sampling effort across Cook Inlet would provide requisite co-occurrence observations of beluga whales and putative prey targets needed to inform foraging patterns for this endangered marine mammal DPS.

The results of the current study emphasize the need for improved quantitative metabarcoding methods in Cook Inlet. We recommend future applications of quantitative eDNA metabarcoding approaches in Cook Inlet leverage qPCR assays for all five salmon species (Chinook, sockeye, coho, pink, and chum; (Elmore et al., 2025) as well as Longfin smelt (Brandl et al., 2015) and Pacific cod (*Gadus macrocephalus*) (Gwak & Nakayama, 2023), comprehensive mock communities of a greater number of putative prey targets for Cook Inlet beluga whales, and sufficient biological and technical replication to enable accurate estimation. In addition, future work could consider applying additional metabarcoding primers to identify gadids like Saffron cod (*Eleginus gracilis*) and walleye pollock (*Gadus chalcogrammus*) that current lack published qPCR assays, but are seasonally abundant (Ledger et al., 2024)). Our findings suggest that future quantitative metabarcoding sampling efforts focussed on foraging patterns during winter months should ensure both longfin smelt and gadids are well characterized.

In addition, we recommend the use of 5 µm filter pore sizes to facilitate easier processing of eDNA samples in Cook Inlet, which has high glacial silt loads, while balancing capture of eDNA sample material (Barnes et al., 2014; Jo & Yamanaka, 2022b, 2022a; Turner et al., n.d.). Utilizing a larger filter would exclude the ∼0.7 µm glacial silt while retaining larger marine mammal sloughed cells and will facilitate easier eDNA collection efforts (Sharma & Burrell, 1970).

Our results suggest that sub-weekly to weekly sampling conducted before, during, and after observations of Cook Inlet beluga whale occupancy in targeted regions would provide sufficient power to identify differential abundances of putative prey targets. We found that temporal variation in eDNA signatures was stronger than spatial (<10 km) or depth (0 to 5 m) variation in this highly tidally influenced system, and recommend allocating sampling effort to a small number of sites at the mouths of estuaries at least 40 km apart, sampled more frequently. Specifically, we recommend targeting key areas of beluga whale habitat utilization to cover seasonally important riverine systems. We also recommend sampling weekly throughout the year where possible, as eDNA sampling across these basins would be particularly useful in informing winter foraging patterns of Cook Inlet beluga whales for which there is limited data on putative prey targets.

Observations from a basin-wide study of Cook Inlet beluga whales and their prey would directly inform future population consequences of disturbance (PCoD) frameworks by providing the requisite observations of the spatial and temporal patterns of putative prey targets (Booth et al., 2020; McHuron et al., 2023). Furthermore, pairing eDNA sampling efforts with passive acoustic data would provide key information needed to understand the interaction between anthropogenic disturbance, prey fields, and Cook Inlet beluga whale behavior. The development of autonomous mooring based eDNA and acoustic samplers could enable such observations at scale, providing uncrewed biodiversity observatories (Yamahara et al., 2025). Generating robust co-located observations of foraging and anthropogenic disturbance are needed to help disentangle cumulative effects for Cook Inlet beluga whales and inform future recovery efforts.

## Conclusion

Our results demonstrate that well calibrated quantitative eDNA sampling on a weekly to sub-weekly frequency paired with visual observations provided sufficient temporal resolution to identify likely prey associations for Cook Inlet beluga whales. These results provide key insights into future basin-wide eDNA-based monitoring of Cook Inlet beluga whale occurrence and prey field patterns to inform future conservation and management of this species.

Ultimately, we demonstrate that quantitative eDNA methods provide a scalable approach for tracking changes in prey fields and marine mammal occurrence, yielding new insight into foraging ecology and supporting broader conservation and recovery efforts. Quantitative eDNA approaches open the door to tracking predator-prey co-occurrence across marine systems and offer a particularly valuable tool for difficult to study species and habitats. In addition, our results highlight the need for careful interpretation of eDNA results, especially in highly dynamic environments with large tides and tidal bores.

## Acknowledgements

This work was supported by the Marine Mammal Laboratory (AFSC, NOAA Fisheries) and NOAA’s Office of Protected Resources. We thank Phil Morin and K. Martien for generously sharing invaluable unpublished marine mammal mitogenomic data with us. Zachary Gold was supported by UW JISAO (Grant #NA15OAR4320063) and UW CICOES (Grant #NA20OAR4320271), NOAA’s Pacific Marine Environmental Laboratory, and NOAA’s Omics Program. We thank Ryan Kelly, Erin D’Agnese, Elizabeth Andruszkiewicz Allan, and Ramon Gallego for guidance and advice in the preparation of this manuscript, data analysis, laboratory processing, and field collection. This manuscript is PMEL contribution number 5882. We thank the many volunteers of the Alaska Beluga Monitoring Program for all their efforts observing Cook Inlet Beluga whales. We are grateful to Alison Gardell and the enthusiastic support of the students and volunteers at University of Alaska Kenai Peninsula College Kachemak Bay Campus, especially Jill Burnham, Courtney Stage, and Teresa Becher. We thank Chris Garner and team at Joint Base Elmendorf-Richardson for conducting Cook Inlet filtration pilot work. We also thank Manuel Castellote for background information and fieldwork coordination. We thank Paul, Barbara, and Laura Crews for graciously hosting Zachary in Girdwood, Alaska during the COVID-19 pandemic during the 2021 field season and providing valuable traditional ecological knowledge of Turnagain Arm.

