## Appendix 1 for "Characterizing the Prey Landscape of Endangered Cook Inlet Beluga Whales Using an eDNA Time Series"

**Appendix 1 Detailed Field and Laboratory Methods**

**Methods**

### **Open Access, FAIR eDNA Methods**

We generated FAIRe Better Biomolecular Ocean Practices (BeBOP) protocols for:

1) eDNA sample collection (<https://doi.org/10.5281/zenodo.24182647>),

2) extraction (<https://doi.org/10.5281/zenodo.24182649>),

3) MiFish 12S PCR amplification (<https://doi.org/10.5281/zenodo.24182633>),

4) Metabarcoding library preparation (<https://doi.org/10.5281/zenodo.24182645>), and

5) Beluga qPCR assay analysis (<https://doi.org/10.5281/zenodo.24182643>)

in accordance with emerging eDNA data management best practices (Samuel et al. 2021; Takahashi et al. 2025; Pitz et al. 2026).

### **eDNA Metabarcoding Methods**

Metabarcoding was performed on all eDNA samples using a previously published two-step PCR library preparation method (Curd et al., 2019; Gold et al., 2021, 2022). First, samples were amplified using the MiFish Universal Teleost primers modified with Illumina Nextera adapter sequences. Specifically, we ran 25 µL PCR reactions consisting of 12.5 µL Qiagen Multiple Mastermix, 1 µL 10 mM Forward primer, 1 µL 10mM Reverse primer, 7.5 µL molecular grade water, and 3 µL of template DNA. Thermocycling used a touchdown PCR protocol described in (Curd et al., 2019): initial denaturation at 95 °C for 15 min, 13 cycles of denaturation at 94 °C for 30 sec, beginning annealing at 69.5 °C for 30 sec (temperature was decreased by 1.5 °C every cycle until 50 °C was reached), and extension at 72 °C for 1 min. 35 additional cycles were carried out at an annealing temperature of 50 °C, followed by a final extension at 72 °C for 10 min. All PCRs were conducted with a non template control and two positive controls comprising tissue extractions from *Oreochromis niloticus* and *Thunnus atlanticus*, two species not native to Cook Inlet, AK. PCR products were run on 2% agarose gels to ensure correct product size.

PCR products were then cleaned using Ampure XP beads (Beckman Coulter, CA, USA) prior to indexing. We prepared sample DNA libraries using indexes from the Nextera Unique Dual Index A through D Kits (Illumina, CA, USA). The PCR was performed using a 25 μL reaction mixture containing 12.5 μL of NEB Phusion High-Fidelity PCR Master Mix (New England Biosystems, MA, USA), 0.625 μL of primer i7, 0.625 μL of primer i5, and 5 ng of PCR 1 template. Thermocycling used the protocol described in Curd et al. 2019: initial denaturation at 95 °C for 3 min, 8 cycles of denaturation at 98 °C for 20 sec, annealing at 56 °C for 30 sec, and extension at 72 °C for 3 min; followed by a final extension at 72 ˚C for 5 minutes. We electrophoresed all indexed PCR products on 2% agarose gels to confirm correct product size. Indexed PCRs were normalized using a SequalPrep Normalization Plate Kit (ThermoFisher, MA, USA) and then pooled in equimolar concentration. The final pool was cleaned and concentrated using Ampure XP beads. Library size was confirmed on an Agilent TapeStation (Agilent, CA, USA). Samples were organized into 5 distinct sequencing runs conducted on a MiSeq using Reagent Kit V3 PE 2x300bp at NWFSC, using loading 6pM of the library with 20% PhiX.
