## Appendix 2 for "Characterizing the Prey Landscape of Endangered Cook Inlet Beluga Whales Using an eDNA Time Series"

**Appendix 2 Detailed Bioinformatics Description**

**Methods**

We used the *Anacapa Toolkit* (Curd et al., 2019) for bioinformatic analysis of eDNA metabarcoding data. The *Anacapa Toolkit* has three modules 1) quality control, 2), amplicon sequence variant parsing, and 3) taxonomic assignment.

The quality control step of the Anacapa Toolkit trims adapter sequences using *cutadapt* (Martin, 2011) and removes low quality reads using FastX-toolkit (Gordon & Hannon, 2010). We used the default parameters in the *Anacapa Toolkit* (All parameters are provided in the github).

The amplicon sequence variant (ASV) parsing step dereplicates reads that pass the quality control module using DADA2 (Callahan et al., 2016). Specifically, we used DADA2 version 1.6 with default parameters to generate ASVs, merge paired reads, and remove chimeric sequences.

Finally, the Anacapa classifier module assigns taxonomy to ASVs using a Bayesian Least Common Ancestor (BLCA) algorithm (Gao et al., 2017) modified to accept alignments from Bowtie 2 (Langmead & Salzberg, 2012) and CRUX reference databases described in (Gold et al., 2021) (https://doi.org/10.5068/D1H963). Briefly, ASVs were aligned to the CRUX reference database using Bowtie2, retaining the top 100 alignments for each ASV. We then conduct multiple sequence alignment for each query ASV to inform a weighted Bayesian posterior probability of taxonomic assignment. Taxonomy is then ultimately assigned based on the lowest common ancestor of the total weighted reference database matches; reliability is evaluated through bootstrap confidence scores which are analogous to percent identity metrics provided by other metabarcoding classifiers (Curd et al., 2019; Gao et al., 2017).Taxonomic assignment was conducted following Gold et al. 2021 using a taxonomic cutoff score of 60 and minimum alignment of 80%.

We note that *Oncorhynchus* salmonid taxonomic assignments were reassigned using a custom script to deal with the high similarity (1-2 bp) in sequence alignments between ASVs. We first removed one reference barcode sequence of chum salmon (*O. keta*) deposited on GenBank (Accession LC091629.1, UW 042256) from Lake Pernatoye, Russia in 1997 with greater similarity to three other *O. nerka* accessions (LC835958.1, OL806600.1, LC340053.1) and appears to be either misidentified in the field, mishandled in the lab, or evidence of hybridization that is not relevant to species identification in Alaska. We then used the DECIPHER R package function clusterize to cluster both ASVs and reference sequences into bins at most 1 bp different from each other (Wright, 2016, 2024). We then accepted the taxonomic assignment provided by Anacapa Toolkit for ASVs and reference sequences that contained the same species level identification within a cluster. For clusters with conflicting species level identifications, ASVs were run through blast against NCBI GenBank. Species level taxonomic assignments were then only retained if all best matches (same percent identity) were to the same species, otherwise we assigned ASVs to the genus level (*Oncorhynchus* spp.).

We then employed a multifaceted post processing approach based on the one developed by Kelly et al. (2018) to remove field contamination, lab contamination, and index hopping artifacts from the metabarcoding data (Gold et al., 2022; McKnight et al., 2019). First, only merged paired reads were retained from the dataset with all forward, reverse, and unmerged paired reads discarded as paired merged reads have the most reliable taxonomic annotations. Second, we evaluated the index hopping by evaluating the proportion of non-target reads that occurred in the two positive control samples. For each ASV, we then subtracted the index hopping proportion as calculated from the positive control from each of the observed sample proportions. Any negative value was set to zero reads. This removed all spurious contaminants that appeared in the positive control samples. Third, we removed known contaminants and off target sequences (e.g. taxonomy assigned to NA, *Homo sapiens;Sus scrofa; Bos taurus;Canis lupis;Gallus gallus,* see code for full list). Fourth, we discarded PCR replicates with less than 20,000 reads and a likelihood of less than 0.01% from a gamma distribution of total sequence read depths. Fifth, we removed low complexity samples where >90% of all reads were assigned to the same species as these are either contaminated or arise from an unexpected process. See the discussion on metabarcode swamping. Sixth, we implemented a site occupancy model as described by Kelly et al. 2018 to retain only ASVs that occurred across multiple sample and technical replicate detections. Functionally, this removed ASVs that were not detected in multiple technical replicates from a given site. Seventh, we removed samples with high dissimilarity between technical PCR replicates which likely arises from a contamination or unintentional laboratory process. Seventh, all reads assigned to the same species were summed, meaning read totals from multiple ASVs assigned to the same species were added together.
