## Appendix 3 for "Characterizing the Prey Landscape of Endangered Cook Inlet Beluga Whales Using an eDNA Time Series"

**Appendix 3. Beluga qPCR Assay Design and Validation**

**Methods**

*Identification of Target Region*

A variable region of the 12S gene region flanked by two conserved regions was selected to design a species-specific qPCR assay based on an alignment of mitogenomes representing 17 cetacean species, including 204 beluga whale sequences (Table S3.1). This variable region supported species-specific probes for *Delphinapterus leucas* targeting a 108 bp amplicon.

The specificity of the primers and probes were tested *in silico* against the consensus sequence of each species using Geneious. The specificity was also tested using the [eDNAssay shiny app](https://nationalgenomicscenter.shinyapps.io/eDNAssay/) (Kronenberger et al., 2022).

**Table S3.1. Marine Mammal Species used to Design Beluga qPCR Assay**

| **Species (Species code)** | **Accessions** | **Count** |
| --- | --- | --- |
| *Balaenoptera acutorostrata* (Bacu) | AP006468, MT410935, NC_005271 | 3 |
| *Balaenoptera musculus* (Bmus) | (Morin et al., 2023), MF409242, NC_001601 | 101 |
| *Berardius bairdii* (Bbai) | NC_005274, Bbai_z0076728_Genewiz_denovo_300821 from P. Morin | 2 |
| *Delphinapterus leucas* (Dleu) | MW316085-MW316288 from Skovrind et al., 2021 | 204 |
| *Eschrichtius robustus* (Erob) | AJ554053, AP006471, MF409244, MF409245, NC_005270 | 5 |
| *Globicephala macrorhynchus* (Gmac) | HM060333 | 1 |
| *Aethalodelphis obliquidens* (Lobl) | MF166863 | 1 |
| *Megaptera novaeangliae* (Mnov) | unpublished full mitogenomes from K. Martien, MF409246, AP006467, and NC_006927 | 44 |
| *Mesoplodon densirostris* (Mden) | P. Morin, 2012 (Morin et al., 2013) | 19 |
| *Mesoplodon europaeus* (Meur) | (Morin et al., 2013) | 8 |
| *Monodon monoceros* (Mmon) | MT251225–MT251289 from Louis et al. 2020 (Louis et al., 2020) | 59 |
| *Orcinus orca* (Oorc) | KF418372–418393, GU187193, KR014267–KR014  271 (Foote et al., 2013; Morin et al., 2015) | 86 |
| *Pseudorca crassidens* (Pcra) | JF289173 | 1 |
| *Phoca vitulina* (Pvit) | NC_001325 | 1 |
| *Stenella longirostris* (Slon) | (Leslie & Morin, 2016) | 103 |
| *Tursiops truncatus* (Ttru) | NC_012059 | 1 |
| *Ziphius cavirostris* (Zcav) | (Morin et al., 2013) | 22 |

*qPCR conditions*

DNA positive controls, DNA of non-target species and eDNA samples as well as non-template controls (distilled water) were quantified using multiplexed TaqMan assays. All amplifications were carried out in triplicate in 10 μl reactions containing 1 × TaqPath ProAmp Multiplex Master Mix (5µL; ThermoFisher Scientific, Waltham, MA US), forward and reverse primers (0.9 μL of 10 µM), hydrolysis probes (0.2 µL of 10 μM), molecular grade water (0.9 µM), internal positive control (IPC) (1 µL of 1,000 copies/µL), and 2 μl of template DNA. Cycling conditions were: initial denaturation step at 95 ̊C for 10 min, followed by 45 cycles of 95 ̊C for 15 s and 60 ̊C for 1 min. Ten-fold serial dilutions starting at ca. 100,000 copies μl^-1^ down to 1 copy μl^-1^ were used to build the standard curve. The above-mentioned tests were analyzed on a QuantStudio 6 (Applied Biosystems). The assays were combined in qPCR multiplexes of up to three different species and included an internal positive control (IPC) to account for PCR inhibition. We used two different IPCs: 1) the previously described *Euryapteryx curtus* broad-billed moa IPC (Ramón-Laca et al., 2021) and 2) the *Hieraaetus moorei* Pouakai (Haast's eagle) IPC (See Tables S3.3). Any delay of more than 0.5 cycles from the IPC at the non-template controls of the PCR was considered inhibition.

**Table S3.2. List of primers used.**

| **Target** | **Primer Name** | **Sequence** | **Length** | **Tm** | **Dye** | **MGB** | **%GC** | **Hairpin Tm** | **Self Dimer Tm** |
| --- | --- | --- | --- | --- | --- | --- | --- | --- | --- |
| Cetaceans | Cet-12S-F | CATGRAGCACGCACACA | 17 | 55.8 - 57.9 |  |  | 56.3 | None | None |
| Cetaceans | Cet-12S-R | CTTGTTACGACTTGTCTC | 18 | 50.1 |  |  | 44.4 | None | None |
| *Delphinapterus leucas* | Dleu-12S-P | TAGATTACTAACCCATGCTAAGCAAGT | 27 | 60.1 | JUN | QSY | 37 | 43.6 | None |
| *Delphinapterus leucas* | Dleu-12S-gblock | TGAATAAGGCCATGAAGCACGCACACACCGCCCGTCACCCTCCTCAAGTAGCATAGCGAAGCCCTAGATTACTAACCCATGCTAAGCAAGTATACGAGAGGAGACAAGTCGTAACAAGGTAAGCGTACCG | 130 | 79.3 |  |  | 50.8 | -11.28 | 256.31 |

**Table S3.3. Internal Positive Control Primers, Probes, and gBlocks.**

| **Target** | **Sequence Name** | **Sequence** | **Length** | **Tm** | **Dye** | **MGB** | **%GC** |
| --- | --- | --- | --- | --- | --- | --- | --- |
| *Euryapteryx curtus* | Moa-IPC-F | CCATCAGACCATACAGTGCAA | 21 | 62.2 |  |  | 47.6 |
| *Euryapteryx curtus* | Moa-IPC-R | GCACTTCAGGTGTATAACTAATGAACG | 27 | 63.7 |  |  | 40.7.4 |
| *Euryapteryx curtus* | Moa-IPC-P | CCGAGATTGTGGGCGCCTGGGTTAC | 25 | 71.7 | VIC | MGB | 64 |
| *Euryapteryx curtus* | Moa-gBlock | GGCTCATGCCCATCAGACCATACAGTGCAATCGGACACGTTTCCCTTGCTCCCCTCTCGTAACCCAGGCGCCCACAATCTCGGGGTGCTTGAAAGCCATACGTTCATTAGTTATACACCTGAAGTGCACA | 130 | 80 |  |  | 53.1 |
| *Hieraaetus moorei* | Haast-IPC-F | CCATATCATCACTATTCATATGTATAC | 27 | 58 |  |  | 29.6 |
| *Hieraaetus moorei* | Haast-IPC-R | TTGAAGATGGGATGACACAAAG | 22 | 60.9 |  |  | 40.9 |
| *Hieraaetus moorei* | Haast-IPC-P | ATGCCCGTATTACATGTATATGGACTAGTACATTA | 35 | 66.9 | ABY | QSY | 34.3 |
| *Hieraaetus moorei* | Haasteagle-IPC-gBlock | CATTCAACTATTTGCCCCATATCATCACTATTCATATGTATACATACTAGTTAATGTACTAGTCCATATACATGTAATACGGGCATGCATCTCTTTGTGTCATCCCATCTTCAAGGACGGGTTTCACTTA | 130 | 79 |  |  | 36.9 |

**Results**

The Dleu-12S assay (forward, primer, and reverse) demonstrated species specificity for beluga both *in silico* and *in vitro* testing (Table S3.4). Only beluga tissues (and not DNA from any other tested cetacean species) amplified in lab testing. Amplification predictions were higher for beluga than all other species with at least 1 mismatch in the probe to all other cetaceans tested, including *Monodon monoceros*, beluga’s closest relative.

**Table S3.4 . Results from eDNAssay tool** for primers Cet-12S-F and Cet-12S-R and the species-specific probe for beluga. ‘Predicted Amplification Score’ is the amplification predictions score obtained with eDNAssay. ‘Mismatches’ indicates the number of mismatches (nucleotides) of the beluga probe against the species detailed in the first column.

|  | **Assay: Dleu-12S-P** | |
| --- | --- | --- |
| **Species** | **Predicted Amplification Score** | **Mismatches** |
| *Balaenoptera acutorostrata* | 0.38 | 10 |
| *Balaenoptera borealis* | 0.43 | 10 |
| *Balaenoptera musculus* | 0.41 | 8 |
| *Balaenoptera physalus* | 0.42 | 9 |
| *Berardius bairdii* | 0.39 | 5 |
| *Delphinapterus leucas* | 0.84 | 0 |
| *Eschrichtius robustus* | 0.47 | 8 |
| *Globicephala macrorhynchus* | 0.69 | 2 |
| *Pseudorca crassidens* | 0.54 | 4 |
| *Megaptera novaeangliae* | 0.42 | 9 |
| *Mesoplodon densirostris* | 0.43 | 8 |
| *Mesoplodon europaeus* | 0.38 | 7 |
| *Lagenorhynchus obliquidens* | 0.62 | 3 |
| *Monodon monoceros* | 0.75 | 1 |
| *Phoca vitulina* | 0.39 | 7 |
| *Tursiops truncatus* | 0.56 | 3 |
| *Orcinus orca* | 0.49 | 4 |
| *Phocoena phocoena* | 0.50 | 3 |
| *Phocoenoides dalli* | 0.50 | 3 |
| *Stenella longirostris* | 0.75 | 2 |
| *Ziphius cavirostris* | 0.38 | 9 |

Following the method of (Klymus et al., 2020), we evaluated the limit of detection of the beluga qPCR assay to be 1 ± 1.06 copies per reaction (median ± sd) with a range from 0.91 to 3.34 copies per reaction (min-max) across all plates analyzed. We found the limit of quantification of the beluga qPCR assay to be 56 ± 2,279.6 copies per reaction (median ± sd) with a range from 8	to 5,139 copies per reaction (min-max) across all plates analyzed. One plate had high variance in observed Ct values resulting in an anonymously high limit of quantification. qPCR assays were only used to track occurrences of Cook Inlet Beluga whales.

**Table S3.5 Limits of Detection and Quantification for Beluga, Eulachon, Coho, and Chinook qPCR Assays.** Units are in copies per reaction.

| **Assay** | **n Plates** | **Min**  **LOD** | **Max**  **LOD** | **Median**  **LOD** | **Mean**  **LOD** | **Sd**  **LOD** | **Min**  **LOQ** | **Max**  **LOQ** | **Median**  **LOQ** | **Mean**  **LOQ** | **Sd**  **LOQ** |
| --- | --- | --- | --- | --- | --- | --- | --- | --- | --- | --- | --- |
| Beluga | 5 | 0.91 | 3.34 | 1.00 | 1.45 | 1.06 | 8.00 | 5139.00 | 56.00 | 1061.20 | 2279.65 |
| Chinook | 5 | 1.12 | 3.94 | 1.71 | 1.99 | 1.12 | 9.67 | 354.00 | 39.00 | 97.57 | 144.68 |
| Coho | 5 | 1.32 | 4.01 | 1.71 | 2.06 | 1.10 | 3.29 | 795.00 | 19.00 | 183.86 | 343.49 |
| Eulachon | 5 | 0.83 | 5.00 | 1.32 | 2.63 | 2.17 | 2.00 | 12.00 | 6.00 | 7.00 | 4.00 |
