## Appendix 4 for "Characterizing the Prey Landscape of Endangered Cook Inlet Beluga Whales Using an eDNA Time Series"

**Appendix 4 Mock Community Results**

**Methods**

We utilized mock communities constructed from vouchered DNA specimens described in (Shelton et al., 2023) and (Gold et al., 2023) to estimate and correct for the known effects of species-specific PCR amplification bias (Shaffer et al., 2025). Here we only used the “Northern” mock communities as they included species relevant to this study (See Shelton et al. 2023 Figure 6).

Specifically, we obtained 22 voucher fish tissue samples from either the Scripps Institute of Oceanography Marine Vertebrate Collection or Northwest Fisheries Science Center Voucher Collection. DNA was then extracted using a Qiagen Blood & Tissue Kit (Qiagen, Inc.), following manufacturer’s protocol. We then amplified a 612 bp fragment of the 12S mtDNA gene that contains the MiFish Universal Teleost 12S primer set (Miya et al. 2015, Collins et al 2021) from each DNA extraction. Long fragment PCR reactions were conducted following (Curd et al., 2019) using Qiagen Multiplex Mastermix and a Touchdown PCR with the exception of using 3 μL starting DNA template for all reactions instead of 1 μL. Nested PCRs were run with 30 cycles. We then quantified the resulting DNA concentration of the PCR products using QuBit Broad Range dsDNA assay (Thermofisher Scientific,Inc.). We then converted the QuBit derived units of mass per volume to copy number per volume using the known fragment length (612bp) and average molecular weight of DNA (650 g/mol). We then constructed the mock communities in both equal and skewed proportions (Table S4.1). The starting concentrations of all six mock communities varied, but were between 120-200 copies/μL. Mock communities were then amplified following the PCR methods described in Appendix 2, identically to our eDNA samples collected in nature.

**Table S4.1 Mock Community Composition**

| **Species** | **Community** | **DNA Ratio** | **DNA Concentration (ng/uL)** | **Total DNA Added (ng)** | **Proportion of DNA** |
| --- | --- | --- | --- | --- | --- |
| *Citharichthys sordidus* | North_Even | 1 | 30 | 30 | 4.5% |
| *Citharichthys stigmaeus* | North_Even | 1 | 30 | 30 | 4.5% |
| *Citharichthys xanthostigma* | North_Even | 1 | 30 | 30 | 4.5% |
| *Engraulis mordax* | North_Even | 1 | 30 | 30 | 4.5% |
| *Sardinops sagax* | North_Even | 1 | 30 | 30 | 4.5% |
| *Sebastes paucispinis* | North_Even | 1 | 30 | 30 | 4.5% |
| *Ophiodon elongatus* | North_Even | 1 | 30 | 30 | 4.5% |
| *Paralabrax clathratus* | North_Even | 1 | 30 | 30 | 4.5% |
| *Paralabrax maculatofasciatus* | North_Even | 1 | 30 | 30 | 4.5% |
| *Paralabrax nebulifer* | North_Even | 1 | 30 | 30 | 4.5% |
| *Scorpaenichthys marmoratus* | North_Even | 1 | 30 | 30 | 4.5% |
| *Bathyraja abyssicola* | North_Even | 1 | 30 | 30 | 4.5% |
| *Hippoglossus stenolepis* | North_Even | 1 | 30 | 30 | 4.5% |
| *Oncorhynchus clarkii* | North_Even | 1 | 30 | 30 | 4.5% |
| *Oncorhynchus gorbuscha* | North_Even | 1 | 30 | 30 | 4.5% |
| *Oncorhynchus keta* | North_Even | 1 | 30 | 30 | 4.5% |
| *Oncorhynchus kisutch* | North_Even | 1 | 30 | 30 | 4.5% |
| *Oncorhynchus mykiss* | North_Even | 1 | 30 | 30 | 4.5% |
| *Oncorhynchus nerka* | North_Even | 1 | 30 | 30 | 4.5% |
| *Oncorhynchus tshawytscha* | North_Even | 1 | 30 | 30 | 4.5% |
| *Squalus acanthias* | North_Even | 1 | 30 | 30 | 4.5% |
| *Thaleichthys pacificus* | North_Even | 1 | 30 | 30 | 4.5% |
| *Hippoglossus stenolepis* | North_Skew_2 | 6 | 20 | 120 | 18.8% |
| *Oncorhynchus clarkii* | North_Skew_2 | 1 | 20 | 20 | 3.1% |
| *Oncorhynchus gorbuscha* | North_Skew_2 | 1 | 20 | 20 | 3.1% |
| *Oncorhynchus keta* | North_Skew_2 | 1 | 20 | 20 | 3.1% |
| *Oncorhynchus kisutch* | North_Skew_2 | 6 | 20 | 120 | 18.8% |
| *Oncorhynchus mykiss* | North_Skew_2 | 1 | 20 | 20 | 3.1% |
| *Oncorhynchus nerka* | North_Skew_2 | 6 | 20 | 120 | 18.8% |
| *Oncorhynchus tshawytscha* | North_Skew_2 | 1 | 20 | 20 | 3.1% |
| *Ophiodon elongatus* | North_Skew_2 | 1 | 20 | 20 | 3.1% |
| *Scorpaenichthys marmoratus* | North_Skew_2 | 1 | 20 | 20 | 3.1% |
| *Squalus acanthias* | North_Skew_2 | 6 | 20 | 120 | 18.8% |
| *Thaleichthys pacificus* | North_Skew_2 | 1 | 20 | 20 | 3.1% |

**Results**

Here we observe amplification bias in our metabarcoding data (observed) versus our mock community (expected) (Figure S4.1).

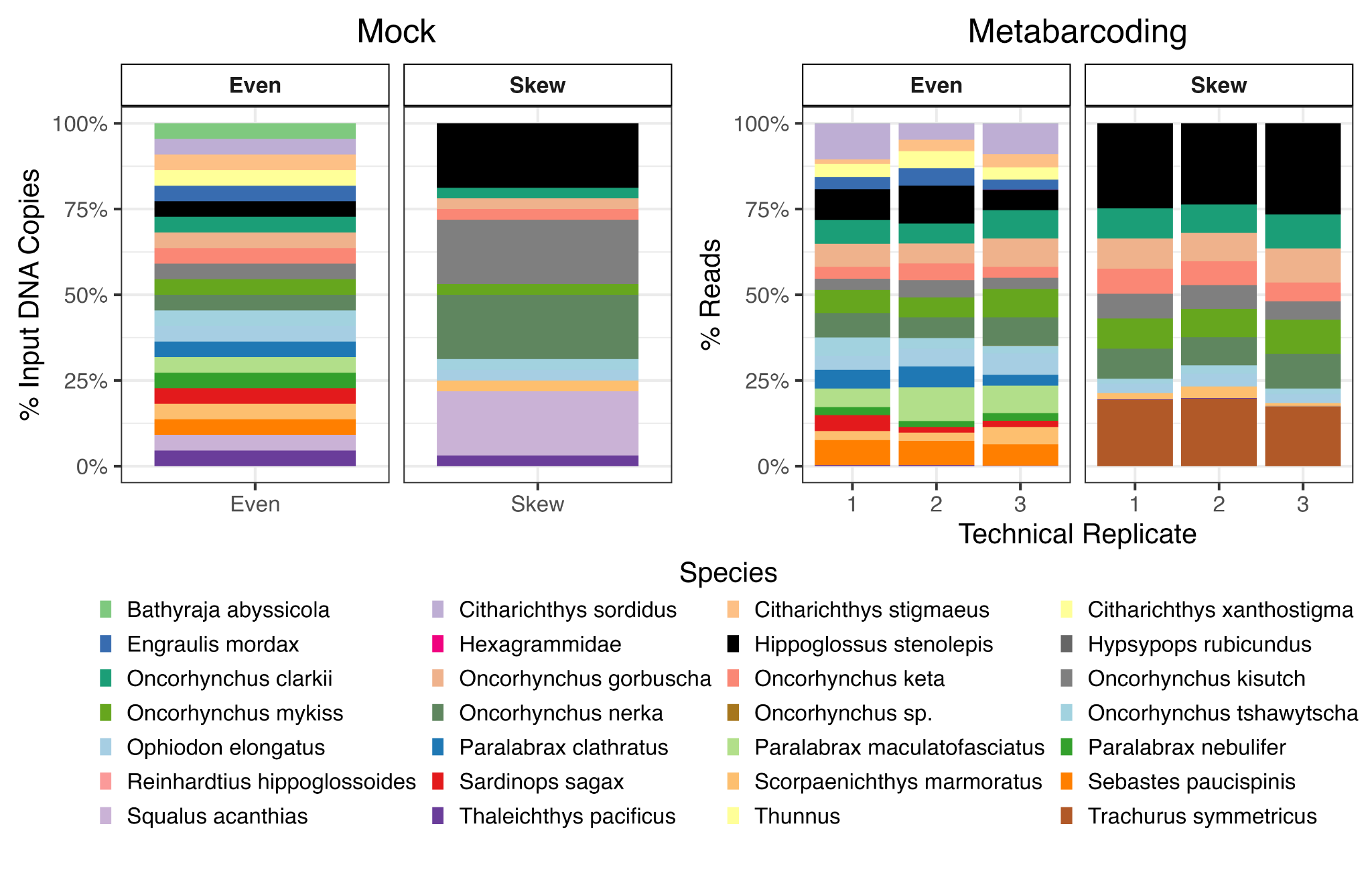

**Figure S4.1 Mock Community Input and Metabarcoding Results.**

We observe biases in metabarcoding results from our known input concentrations. In particular, Chondrichthyans (*Bathyraja abyssicola* and *Squalus acanthias*) are severely under-represented in the metabarcoding results. We observed variability across technical replicates.

We focus our analyses on five species of salmon Pink (*Oncorhynchus gorbuscha*), Chum (*O. keta*), Coho (*O. kisutch*), Sockeye (*O. nerka*), and Chinook (*O. tshawytscha*) and observe patterns of amplification bias across taxa albeit subtler than for Chondrichthyan taxa. Given the compositional nature of metabarcoding we analyzed our mock communities using the additive log ratio transformation. Using this transformation, we designated Chinook as our reference species (Amplification Efficiency = 0; Table S4.1). We removed all other species from our mock community analyses and recalculated total reads as just those assigned to these five *Oncorhynchus* species. We then recomputed corresponding proportional reads.

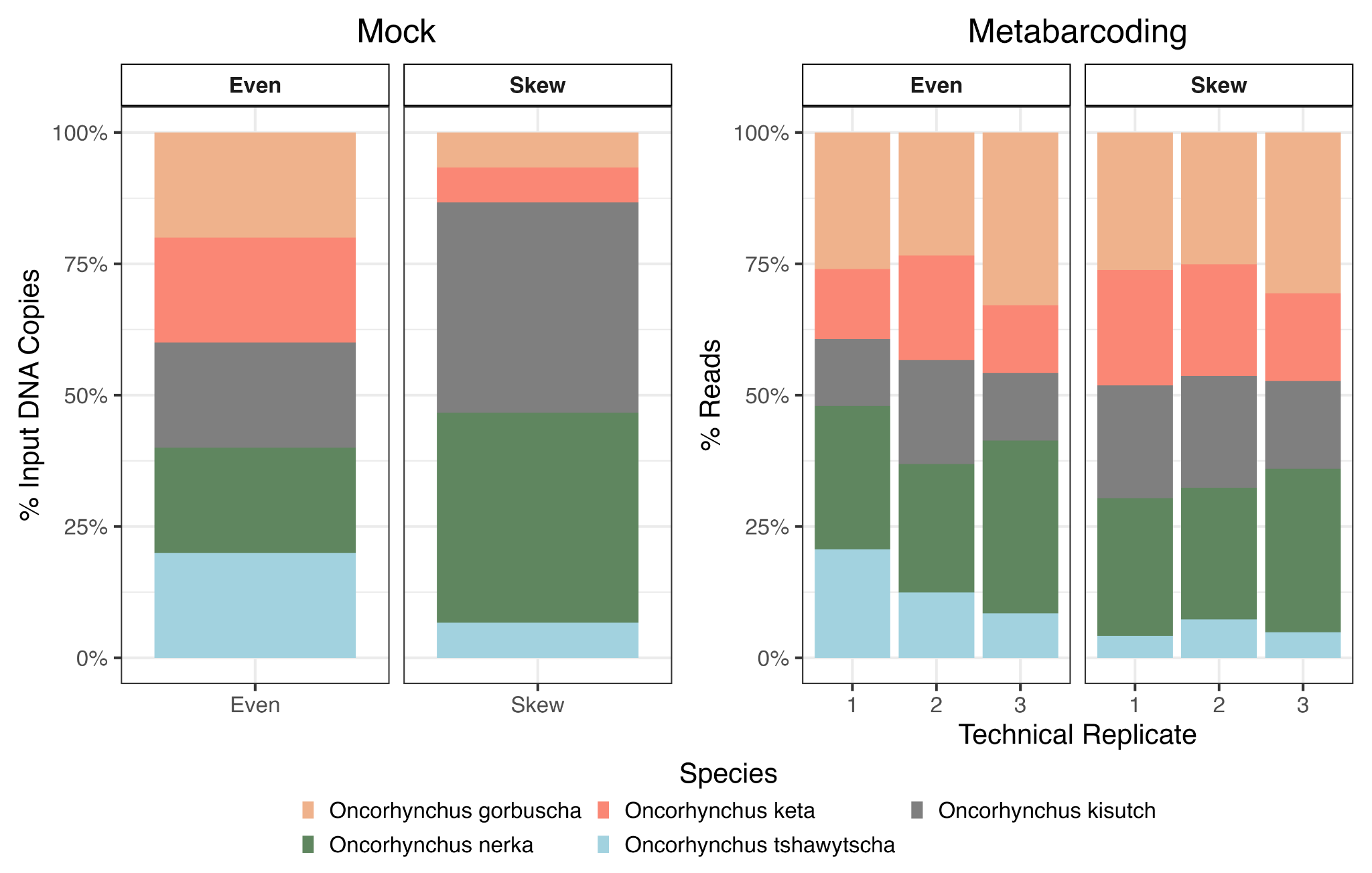

**Figure S4.2 Salmon Mock Community Input and Metabarcoding Results.**

We observe biases in metabarcoding results from our known input concentrations for salmon taxa.

The further the amplification efficiency (additive log ratio) is from zero, the greater the amplification bias. We then converted the amplification efficiency values into a multiplicate bias factor accounting for the 39 PCR cycles employed to provide a more easily understandable metric. Here we see the *O. gorbuscha* and *O. keta* dramatically overamplify compared to *O. kisutch.*

**Table S4.2 Estimated Amplification Efficiency Bias and Observed Mean Relative Error.**

| **Species** | **Amplification Efficiency (ALR)** | **Multiplicative bias factor** | **Result** |
| --- | --- | --- | --- |
| *Oncorhynchus gorbuscha* | 0.030 | 3.28 | Over amplifies |
| *Oncorhynchus keta* | 0.019 | 2.06 | Over amplifies |
| *Oncorhynchus kisutch* | -0.005 | 0.840 | Under amplifies |
| *Oncorhynchus nerka* | 0.008 | 1.36 | Over amplifies |
| *Oncorhynchus tshawytscha* | 0.000 | 1.0 | Reference Species |

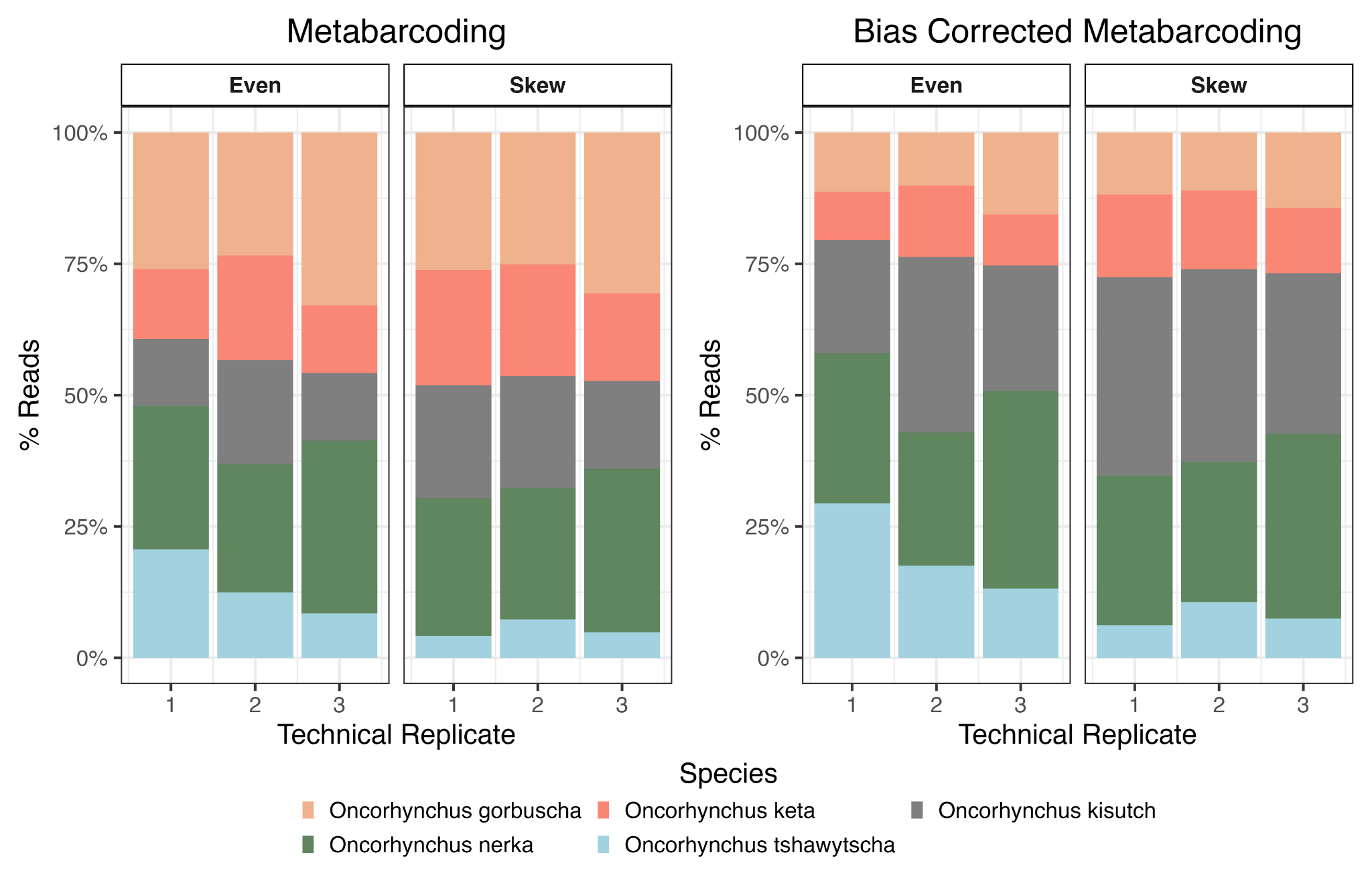

**Figure S4.3 Salmon Bias Corrected Metabarcoding Results.**

Bias corrected reads better approximate known inputs into the mock communities, particularly the reduction in *O. gorbuscha* and *O. keta* proportions and increase in *O. kisutch.*

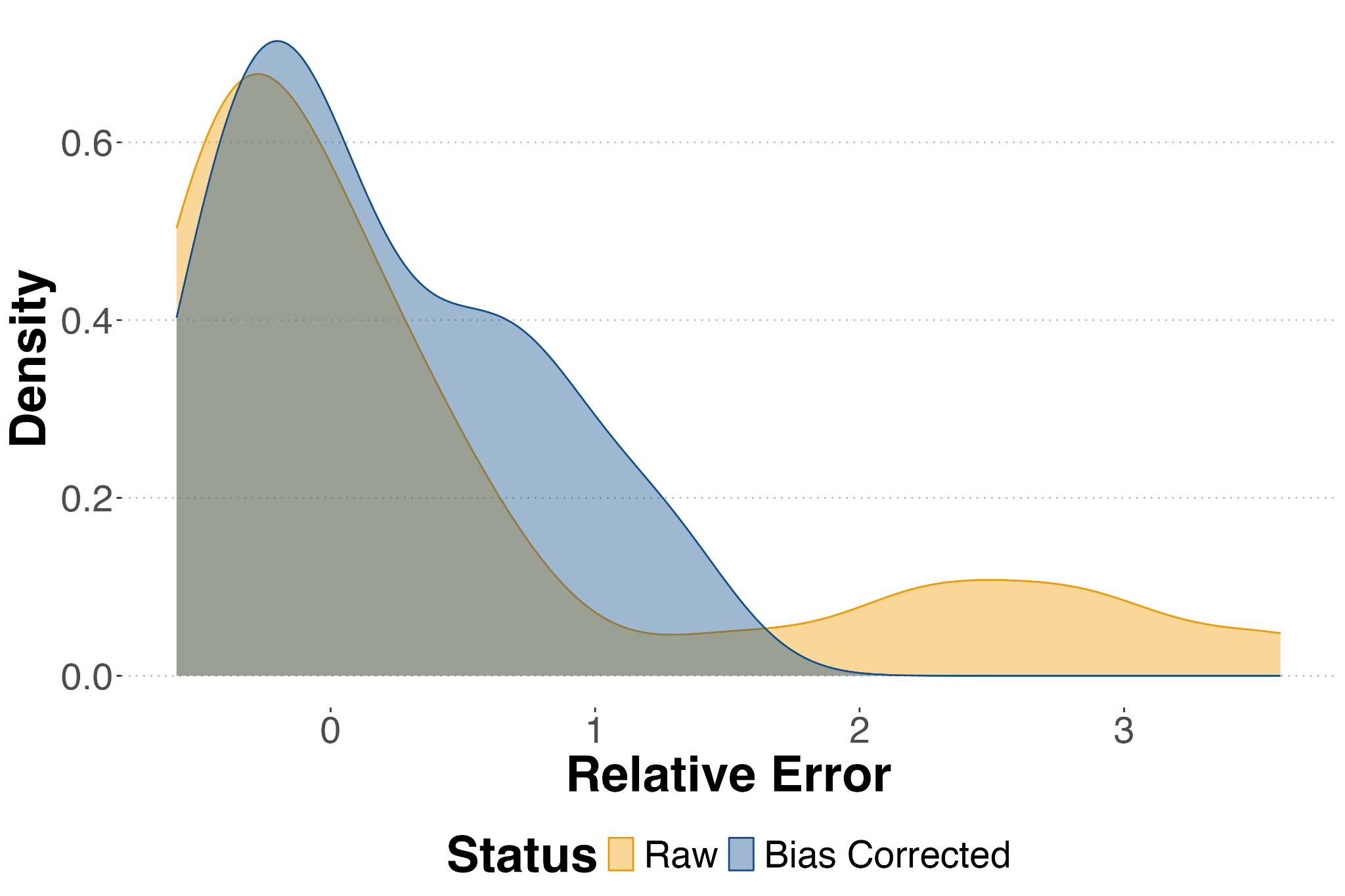

**Figure S4.4 Bias Corrected Mock Communities Have Decreased Relative Error.**

Bias corrected reads better approximate known inputs into the mock communities with reduced relative error*.* Relative error was calculated as (Observed Proportions - Expected Proportions) / Expected Proportions.

Applying the log-linear bias correction resulted in a lower Aitchison distance to the known true composition compared to the raw observed proportions. Mean error for raw metabarcoding reads was 1.36 ± 0.63 (mean ± sd) while mean error for biased corrected metabarcoding reads was 0.99 ± 0.06. We note distances between bias corrected and raw reads were not significantly different (Wilcoxon signed-rank test, p = 0.15). However, this was largely driven by the small sample size (n=6 mocks), limiting statistical power. Mean distances, particularly large relative errors were consistently reduced.

The resulting amplification efficiencies were then used to bias correct our observed metabarcoding reads, providing a more accurate estimate of underlying proportional DNA abundance used with qPCR data to derive absolute DNA concentration abundance estimates for the five species of salmon analyzed here in.
