## Appendix 5 for "Characterizing the Prey Landscape of Endangered Cook Inlet Beluga Whales Using an eDNA Time Series"

**Appendix 5 qPCR Anchor Comparisons**

**Results**

Here we compared how well our quantitative metabarcoding model performed when using Chinook (*O. tshawytscha)* or Coho *(O. kisutch)* as the anchor species. Here we define the anchor species as the species used to estimate the total DNA concentrations per sample that was observed by both metabarcoding and qPCR approaches. Total DNA concentration in a given sample is estimated by dividing the total observed DNA concentration for the anchor species by the proportion of that species derived from bias corrected metabarcoding.

We first compared predicted total DNA concentrations estimated for each sample when using Coho (*O. kisutch*) and Chinook (Figure S5.1). Samples with less than 5% read abundance of either species had poor estimates of total DNA. When excluding samples with less than 5% read abundance of either species, we found a weak correlation between estimated total DNA concentrations (R2 =0.28, p <0.001). Total DNA concentrations entirely fell below the 1:1 line indicating that Coho derived total DNA concentration estimates were systematically higher than Chinook derived estimates.

However, when comparing predicted DNA concentrations on a per species per bottle basis, we found a strong correlation between reads predicted from Coho and Chinook anchors (R2 = 0.85, p <0.0001; Figure S5.2). Samples generally fell along the 1:1 line suggesting per species per bottle estimates of DNA concentrations were more similar than total DNA concentration estimates.

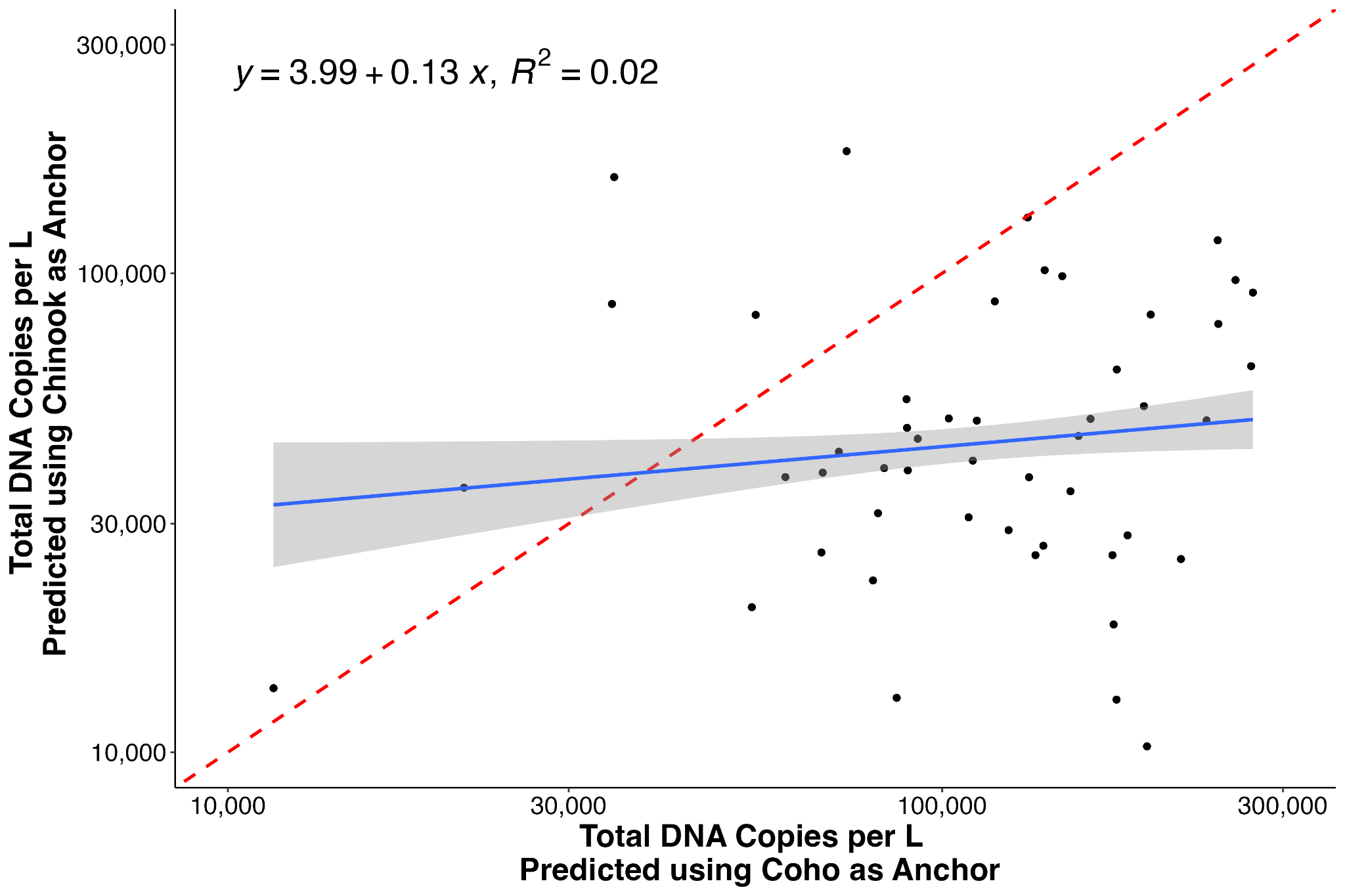

**Figure S5.1.** **Comparison of Total DNA Copies per L Predicted from Coho and Chinook as Anchor Species.** We find a weak positive correlation between total DNA copies per L predicted from Chinook and Coho anchors. We see a bias towards higher total DNA concentrations from Coho. The 1:1 line is plotted in red.

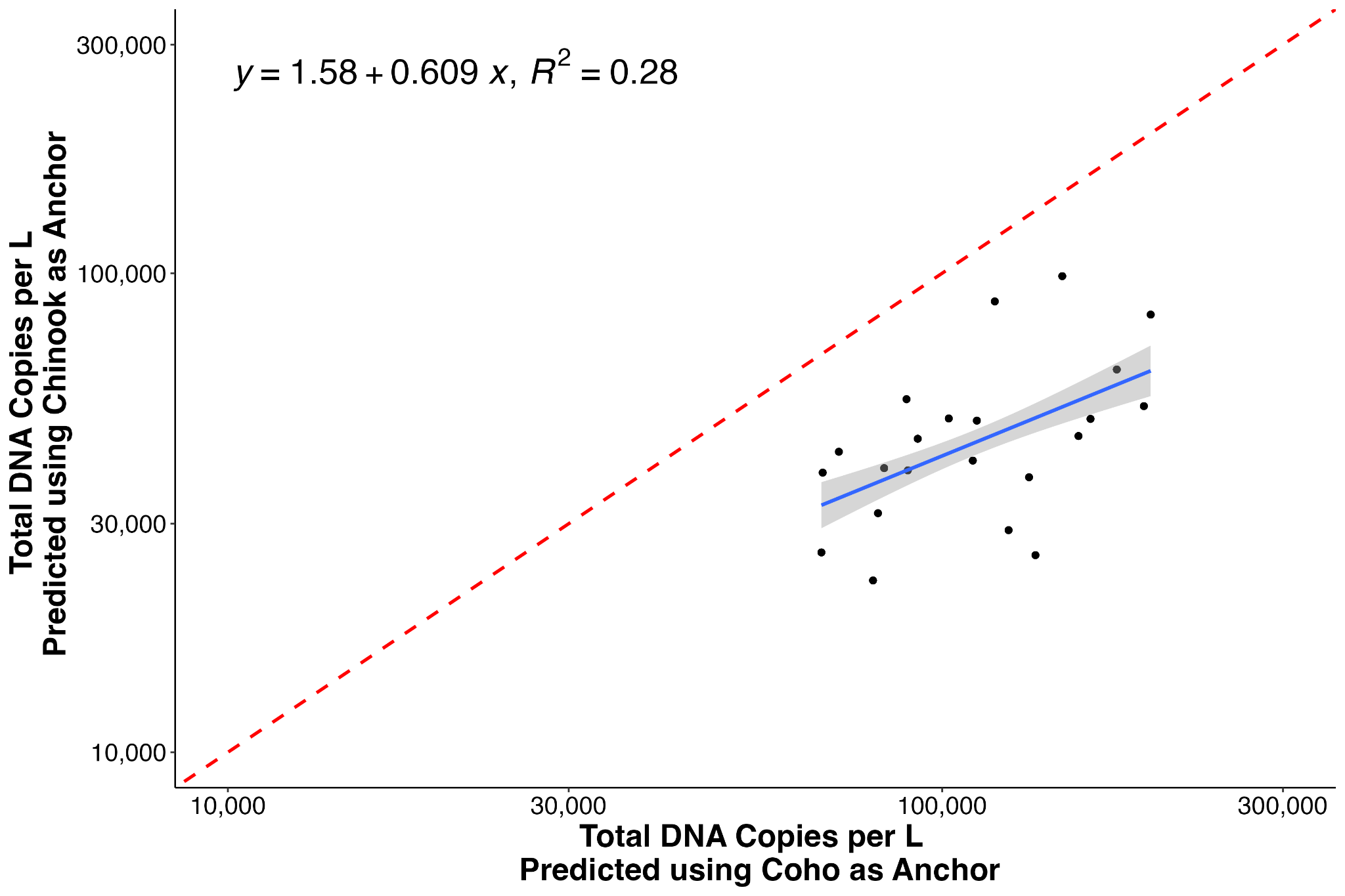

**Figure S5.2.** **Comparison of Total DNA Copies per L Predicted from Coho and Chinook as Anchor Species with Samples with Low Respective Abundance Excluded.** We find a weak positive correlation between total DNA copies per L predicted from Chinook and Coho anchors. We see a bias towards higher total DNA concentrations from Coho. The 1:1 line is plotted in red. Samples with less than 5% read abundance for either species were excluded.

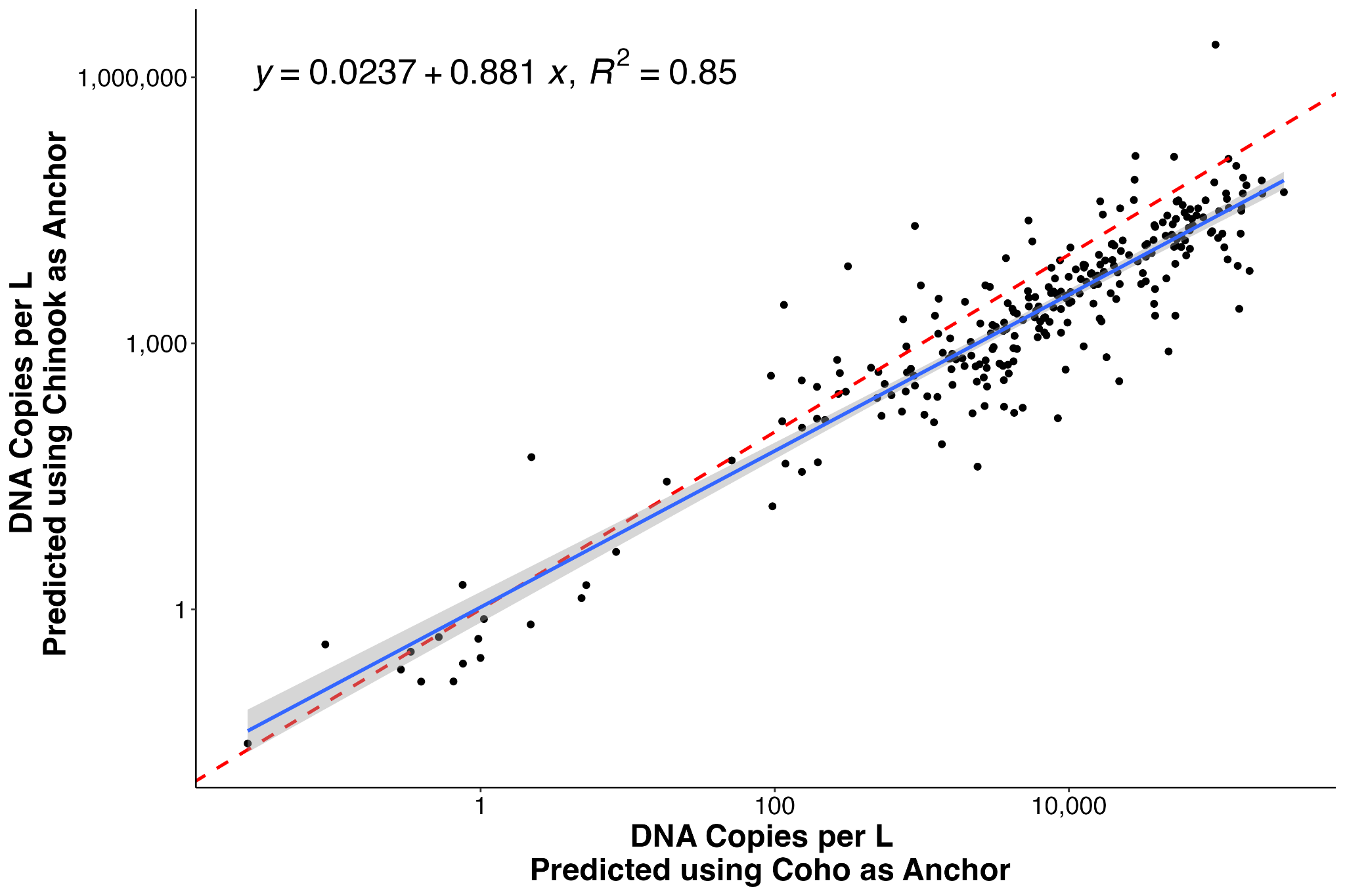

**Figure S5.3.** **Per Bottle Per Species Comparison of DNA Copies per L Predicted from Coho and Chinook as Anchor Species.** We find a strong positive correlation between predicted DNA copies per liter per bottle per species. The 1:1 line is plotted in red. All samples were included.

Furthermore, we similarly observe nearly identical temporal patterns in predicted Chinook and Coho DNA concentrations in Twenty Mile River from both anchor species (Figures S5.4-5). We observed similar patterns of higher Coho DNA concentrations during the period of beluga occupancy in Twentymile River and lower concentrations of Chinook DNA. These results highlight the robustness of our findings across anchor species.

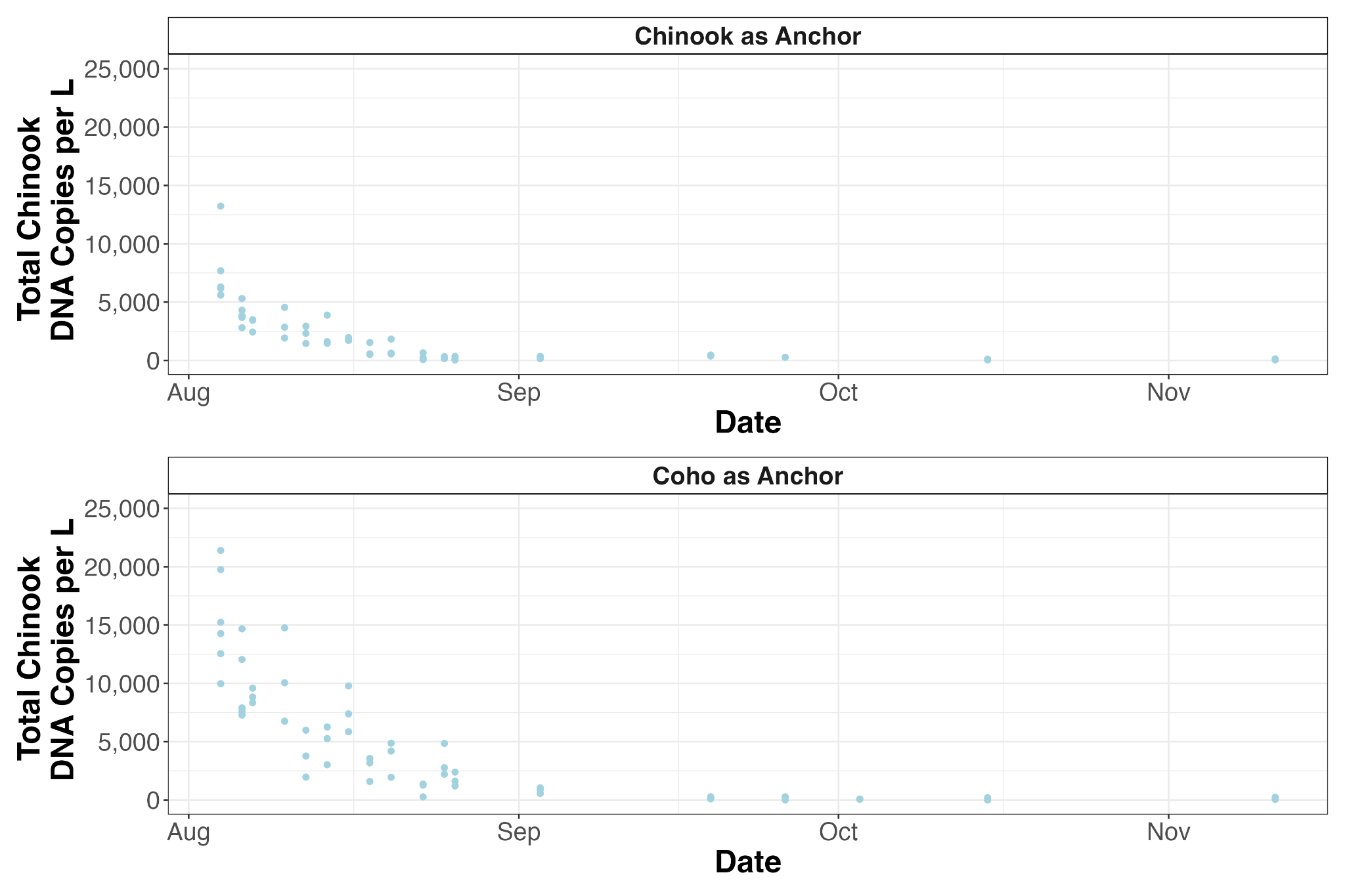

**Figure S5.4.** **Comparison of estimated total Chinook concentrations from Coho and Chinook anchors in Twentymile River.** We observe a similar decline in total Chinook DNA concentrations in Twentymile River regardless of choice in anchor species.

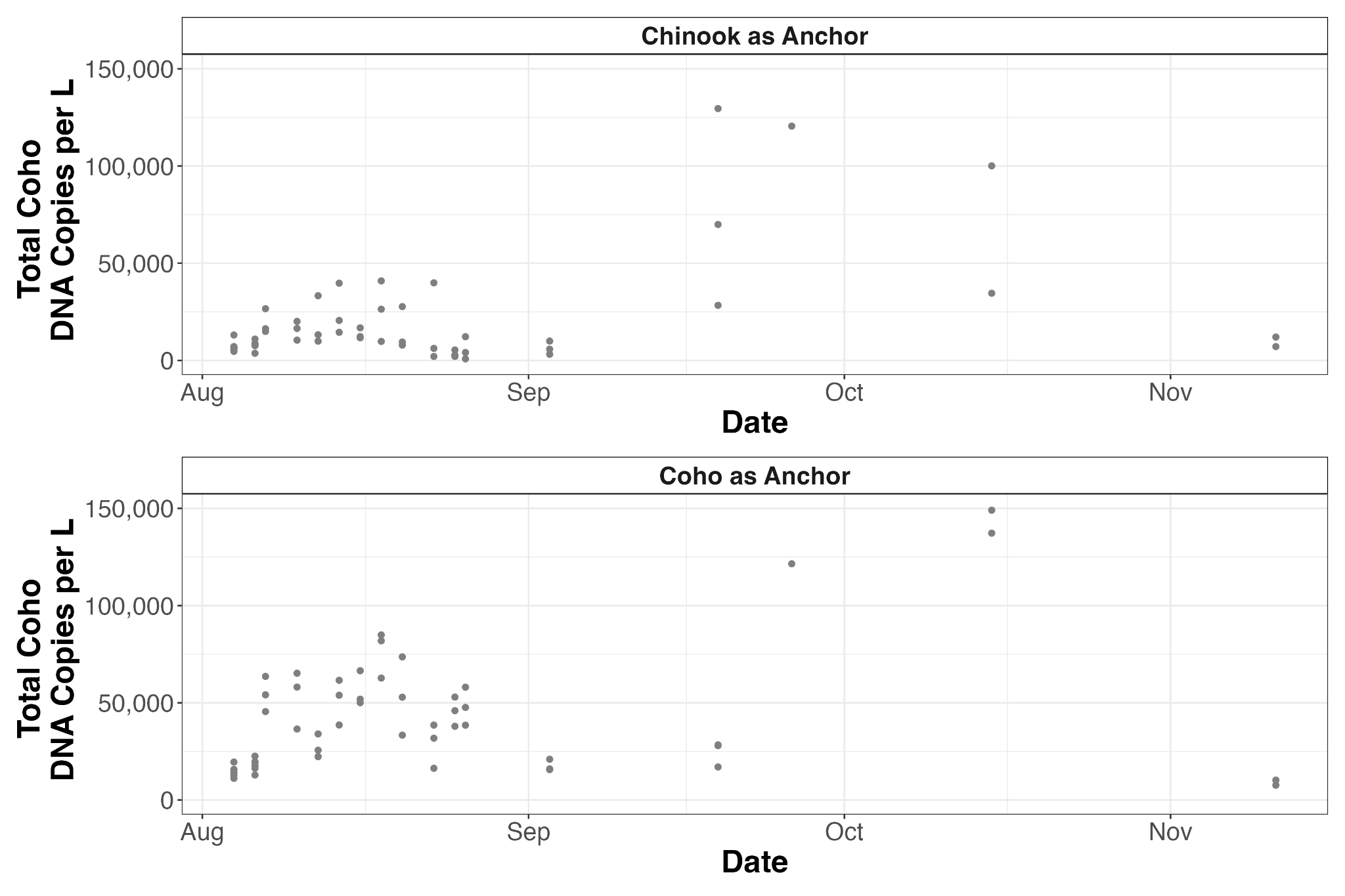

**Figure S5.5.** **Comparison of estimated total Coho concentrations from Coho and Chinook anchors in Twentymile River.** We observe a similar pattern in total Coho DNA concentrations in Twentymile River regardless of choice in anchor species. We note that when using Coho as an anchor, we generally predicted higher concentrations of Coho than when we used Chinook as an anchor with the exception of mid September sampling time pint.

However, we do note that we did not find significant differences in Coho DNA before and during beluga occupancy in Twentymile River when using Chinook as the anchor species (Figure S5.6). We did find significant differences in Chinook and Pink DNA before and during beluga occupancy when using Chinook as the anchor species coherent with results presented in the main manuscript using Coho as the anchor species.

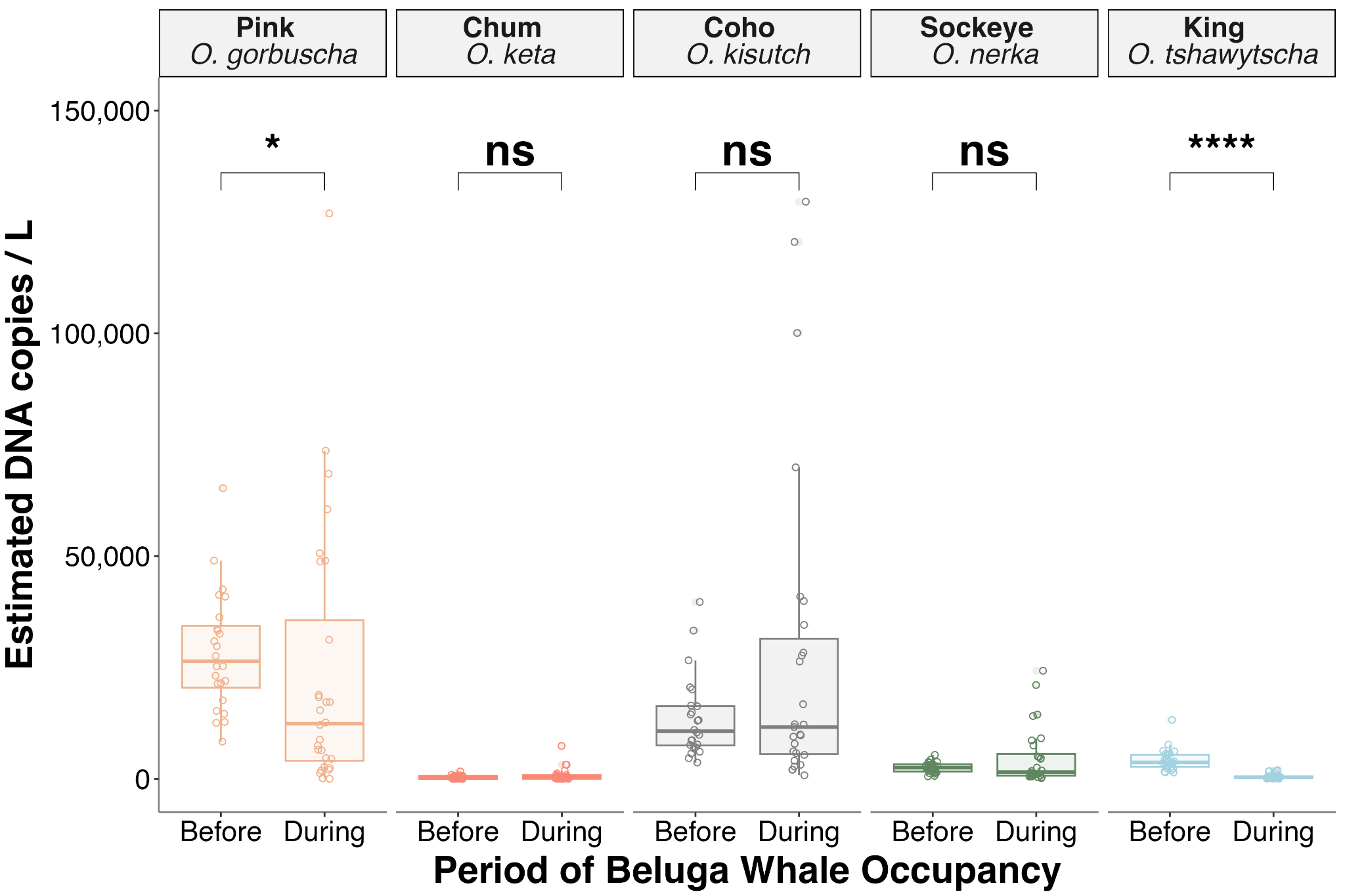

**Figure S5.6 Quantitative Metabarcoding Derived Salmon Abundance Concentrations Over Time in Twentymile River using Chinook as the Anchor Species.** Boxplots represent median, interquartile range, and whiskers extend to maximum and minimum values. Significance values derived from pairwise Kruskal-Wallis comparisons of estimated DNA abundance are shown (ns: p > 0.05, *: p <= 0.05, **: p <= 0.01, ***: p <= 0.001, ****: p <= 0.0001).

Partial alignment in cross anchor comparisons highlights a challenge of the employed quantitative metabarcoding approach and the need to utilize an appropriate anchor species that is sufficiently abundant and present across all datasets.

These results are in part explained by differences in prevalence and abundance of both species throughout the dataset. In Twentymile River, Coho were more frequently detected (Metabarcoding: total detections = 62 out of 62, qPCR: total detections = 62 out of 62) than Chinkook (Metabarcoding: total detections = 60 out of 62, qPCR: total detections = 58 out of 62). Furthermore, Coho was more proportionally abundant in the Twentymile River metabarcoding dataset with a mean of 37.1% ± 25.8% reads (max = 98.6%, min = 10.2 %) across all samples compared to a mean of 4.6% ± 4.7% (max=15.9%, min= 2.2*10^-5). Specifically, Coho and Chinook exhibited opposite patterns in abundance in both the qPCR and metabarcoding datasets with Coho increasing in absolute and proportional abundance when Chinook was rarest and least abundant. When using Chinook as the anchor species to estimate Coho during the period of beluga occupancy, this required dividing relatively lower concentration Chinook qPCR observations (<500 copies per L) by proportionally rare Chinook bias corrected metabarcoding reads (<5%) to estimate Coho DNA concentrations when it was maximally abundant (> 70% read abundance, > 10,000 copies per L). This resulted in high variance of estimated Coho DNA concentrations during the period of beluga occupancy in Twentymile River.

Previous work has shown that variance in both qPCR and metabarcoding data is a function of underlying concentration (Shelton et al., 2026). Thus, using a rarer and less abundant anchor species to estimate a more abundant taxa would result in more variable, underestimates of DNA concentrations. Our results align with previous simulation and observational data (Allan et al., 2023; Gold et al., 2023; Shaffer et al., 2025; Shelton et al., 2022). Thus we argue to best estimate quantitative abundance, an ideal anchor species would be abundant in all samples.

Fortunately for us, Coho was present in all Twentymile River samples with at least 10% read abundance, making it a far more ideal candidate as an anchor species than Chinook which was proportionally rare in most of the time series. Our results highlight that our approach is fairly robust to anchor species selection, especially when anchor species are not proportionally rare in a sample (< 5% abundance). Thus, given the need for a prevalent and abundant anchor species we used Coho as the anchor species in our analyses.

Together, these results lend credence to our mechanistic model of quantitative metabarcoding shared here and highlight the importance of selecting an anchor species with high prevalence across all samples to anchor proportional metabarcoding data and propagate estimated absolute DNA concentrations.

Allan, E. A., Kelly, R. P., D’Agnese, E. R., Garber-Yonts, M. N., Shaffer, M. R., Gold, Z. J., & Shelton, A. O. (2023). Quantifying impacts of an environmental intervention using environmental DNA. *Ecological Applications: A Publication of the Ecological Society of America*, *33*(8), e2914.

Gold, Z., Shelton, A. O., Casendino, H. R., Duprey, J., Gallego, R., Van Cise, A., Fisher, M., Jensen, A. J., D’Agnese, E., Andruszkiewicz Allan, E., Ramón-Laca, A., Garber-Yonts, M., Labare, M., Parsons, K. M., & Kelly, R. P. (2023). Signal and noise in metabarcoding data. *PloS One*, *18*(5), e0285674.

Shaffer, M. R., Andruszkiewicz Allan, E., Van Cise, A. M., Parsons, K. M., Shelton, A. O., & Kelly, R. P. (2025). Observation bias in metabarcoding. *Molecular Ecology Resources*, *25*(7), e14119.

Shelton, A. O., Kachel, S. M., Shaffer, M. R., Brandão-Dias, P. F. P., Jacobson, E. K., Thomas, L., & Kelly, R. P. (2026). Estimating absolute DNA abundance from replicated metabarcoding. *Methods in Ecology and Evolution*, (2041-210x.70427). https://doi.org/10.1111/2041-210x.70427

Shelton, A. O., Ramón-Laca, A., Wells, A., Clemons, J., Chu, D., Feist, B. E., Kelly, R. P., Parker-Stetter, S. L., Thomas, R., Nichols, K. M., & Park, L. (2022). Environmental DNA provides quantitative estimates of Pacific hake abundance and distribution in the open ocean. *Proceedings. Biological Sciences*, *289*(1971), 20212613.
