## Appendix 6 for "Characterizing the Prey Landscape of Endangered Cook Inlet Beluga Whales Using an eDNA Time Series"

**Appendix 6 Supplemental qPCR and Metabarcoding Figures**

**Results**

**Twentymile River**

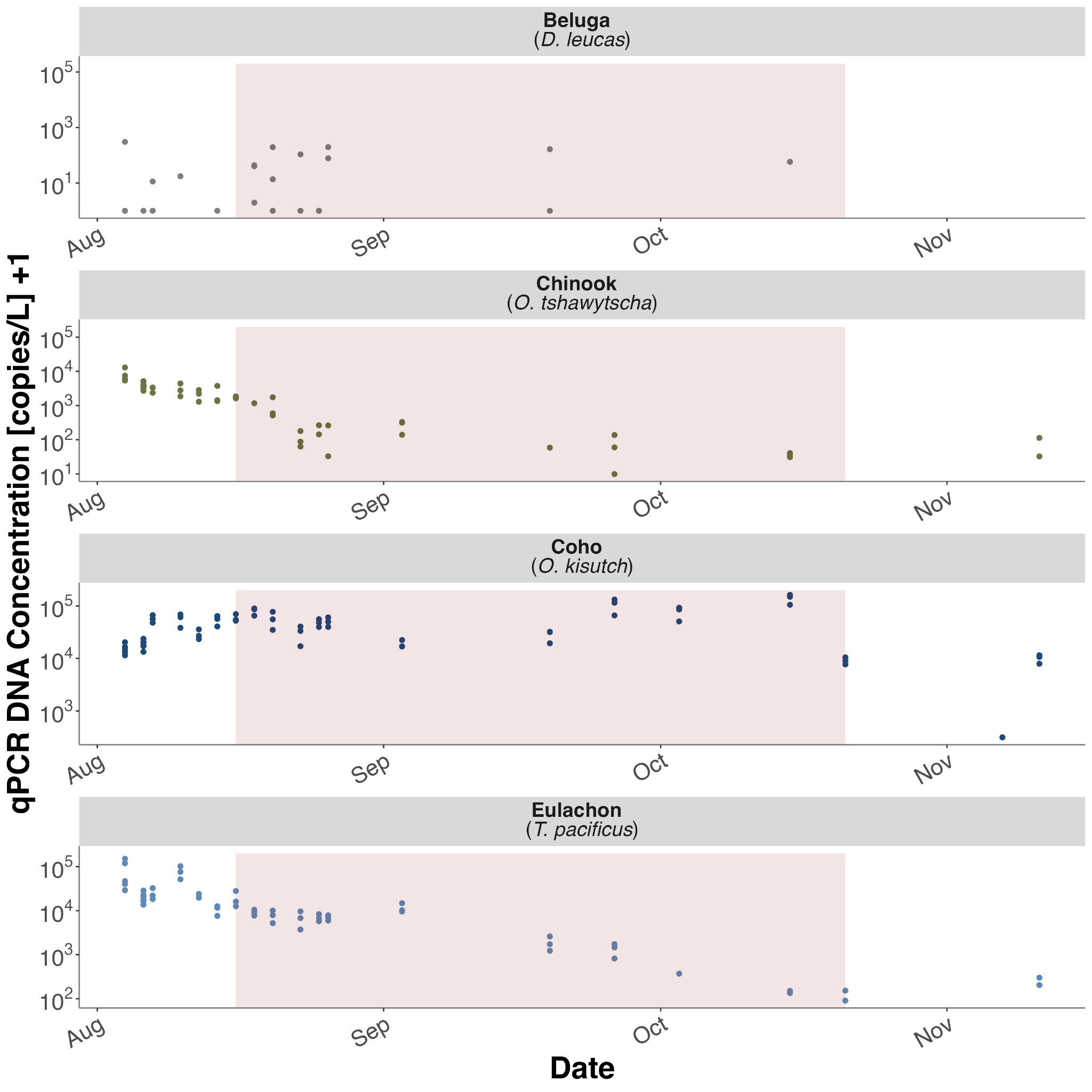

**Figure S6.1 Twentymile River qPCR Results Over Time.**

qPCR derived DNA concentrations in Twentymile River. Only detections are plotted. Note Y axis is logarithmic and limits are scaled per species to highlight intraspecific patterns. Period of beluga occupancy is shaded in red.

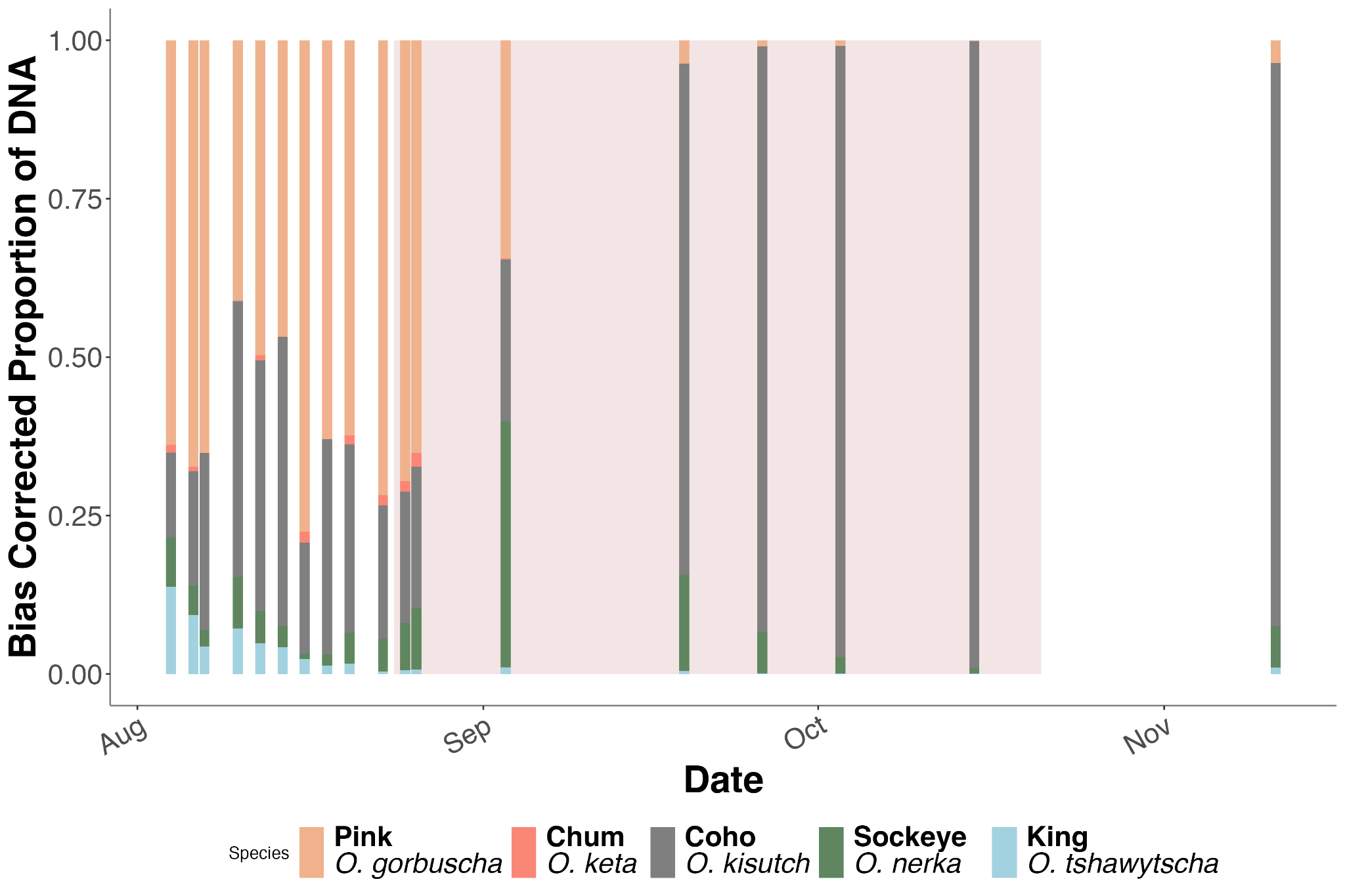

**Figure S6.2 Twentymile River Bias Corrected Metabarcoding Results Over Time.**

Stacked bar plots of bias corrected metabarcoding proportions. Species are color coded. Period of beluga occupancy is shaded in red. Only these five salmon species were analyzed for quantitative metabarcoding.

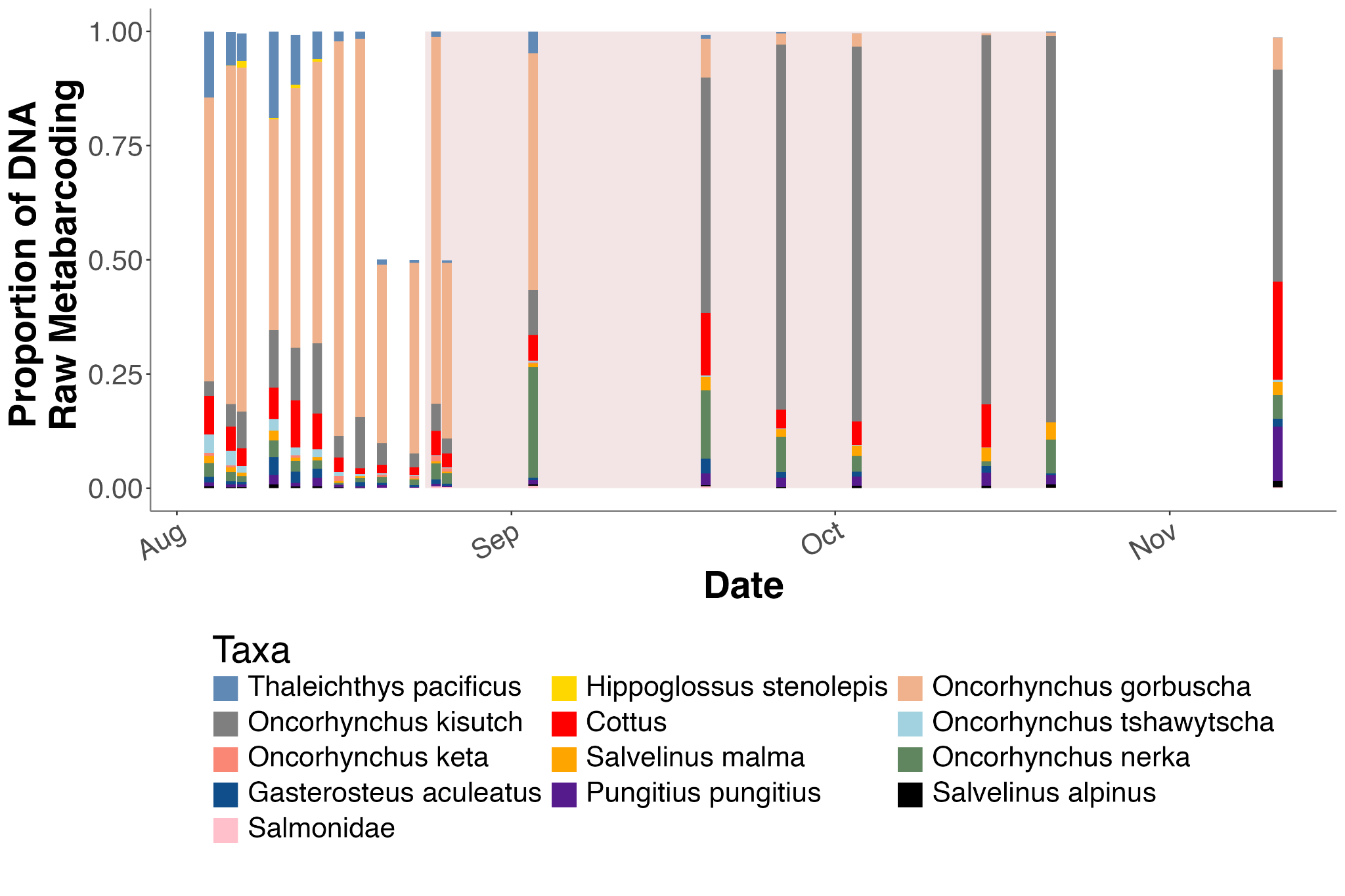

**Figure S6.3 Twentymile River Raw Metabarcoding Results Over Time.**

Stacked bar plots of raw metabarcoding proportions. Taxa are color coded. Only bony fish species with at least 5 detections and 5,000 reads are shown. Period of beluga occupancy is shaded in red.

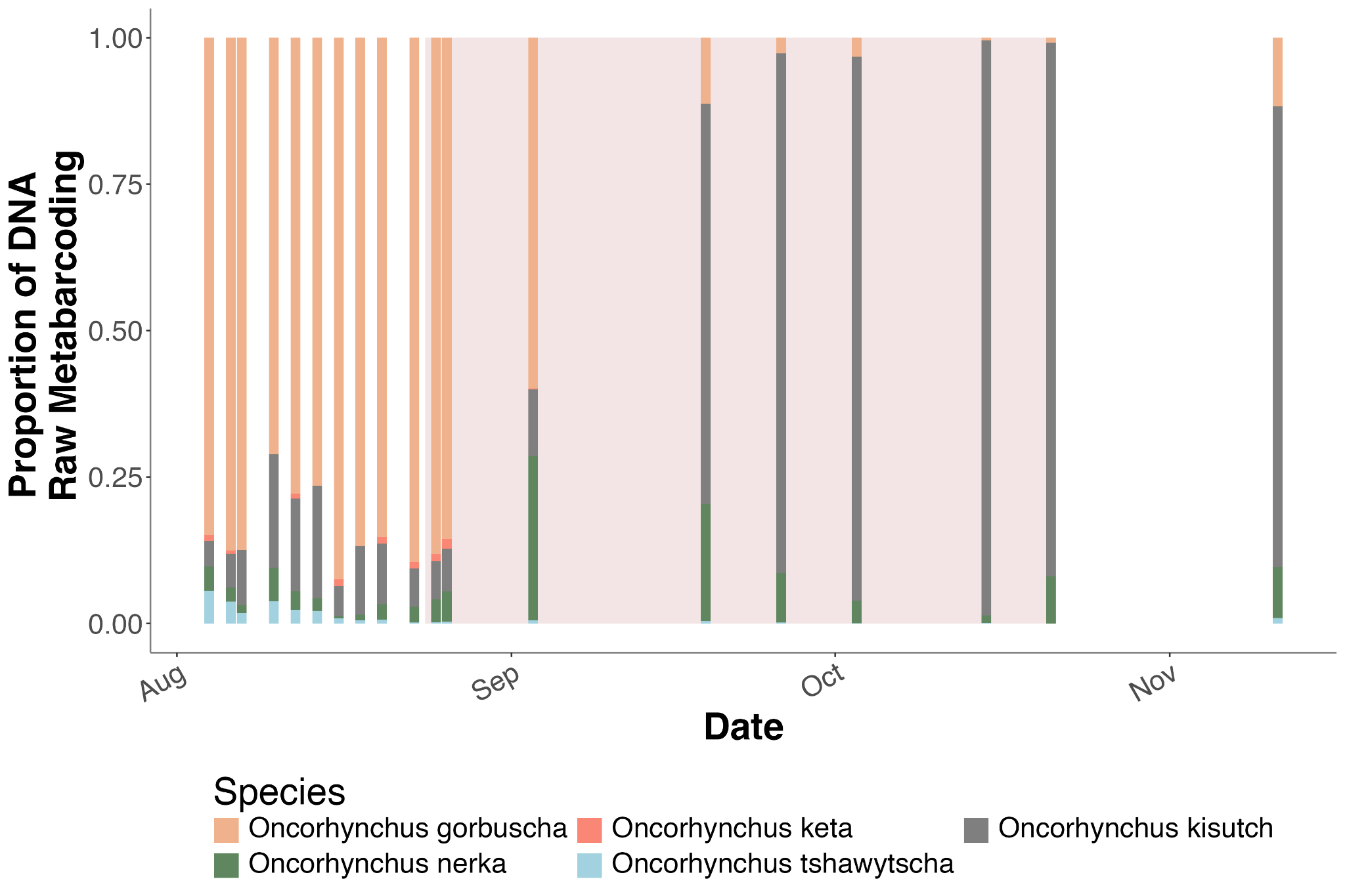

**Figure S6.4 Twentymile River Raw Salmon Metabarcoding Results Over Time.**

Stacked bar plots of raw metabarcoding proportions. Taxa are color coded. Only the five salmon species analyzed for quantitative metabarcoding are shown. Period of beluga occupancy is shaded in red.

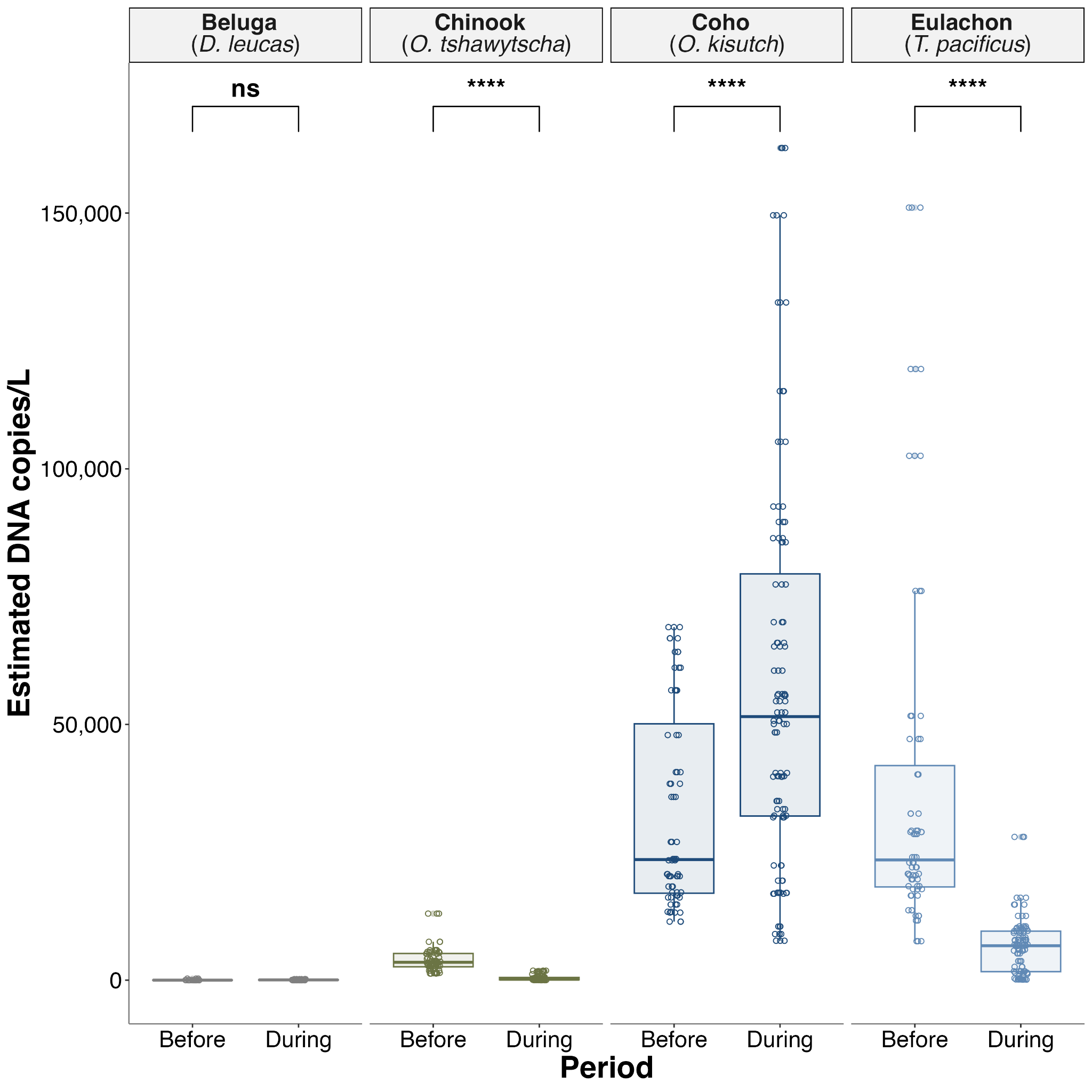

**Figure S6.5 qPCR Derived Salmon Abundance Concentrations Over Time in Twentymile River.** Boxplots represent median, interquartile range, and whiskers extend to maximum and minimum values. Significance values derived from pairwise Kruskal-Wallis comparisons of estimated DNA abundance are shown (ns: p > 0.05, *: p <= 0.05, **: p <= 0.01, ***: p <= 0.001, ****: p <= 0.0001).

**Kenai River**

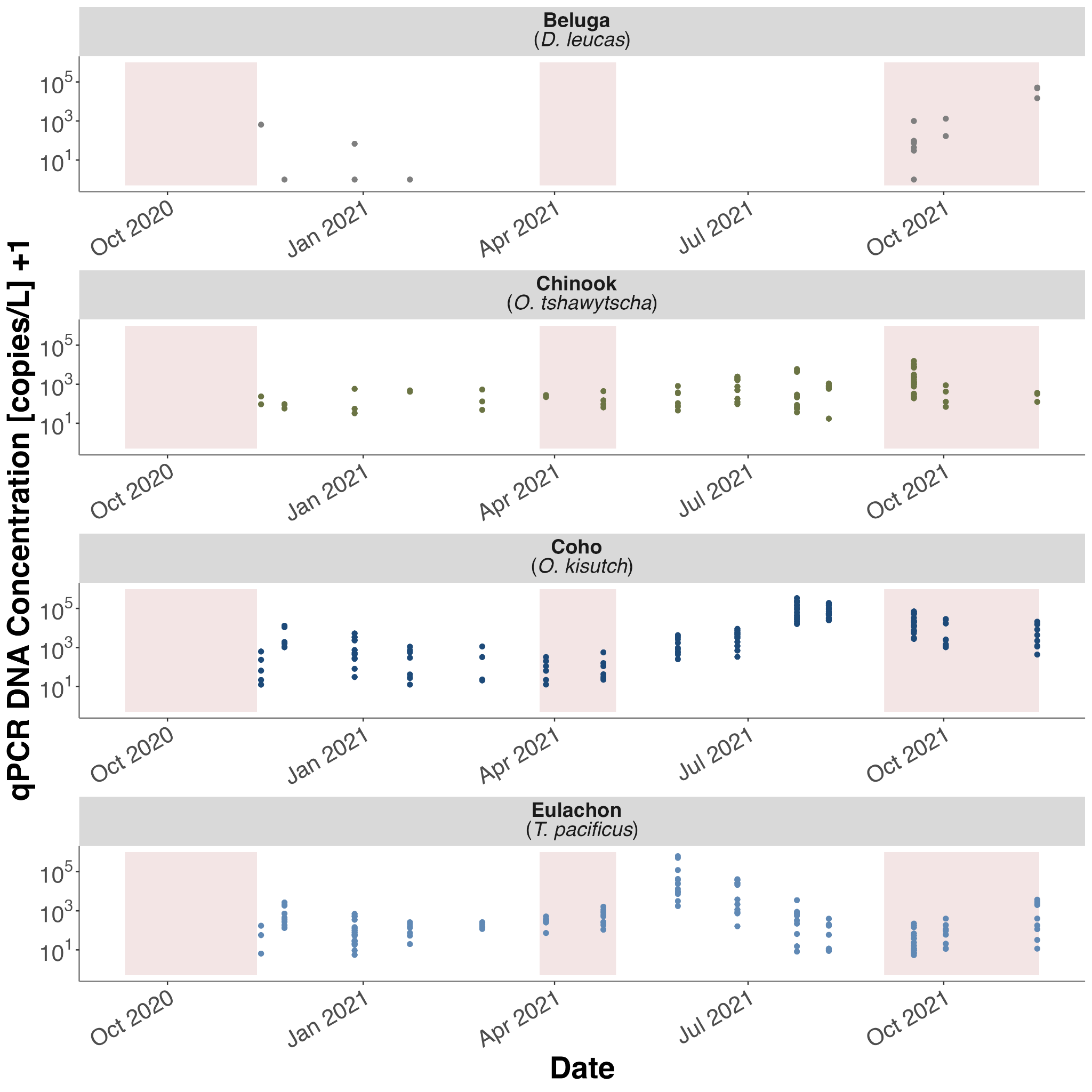

**Figure S6.6 Kenai River qPCR Results Over Time.**

qPCR derived DNA concentrations in Twentymile River. Only detections are plotted. Note Y axis is logarithmic and limits are scaled per species to highlight intraspecific patterns.

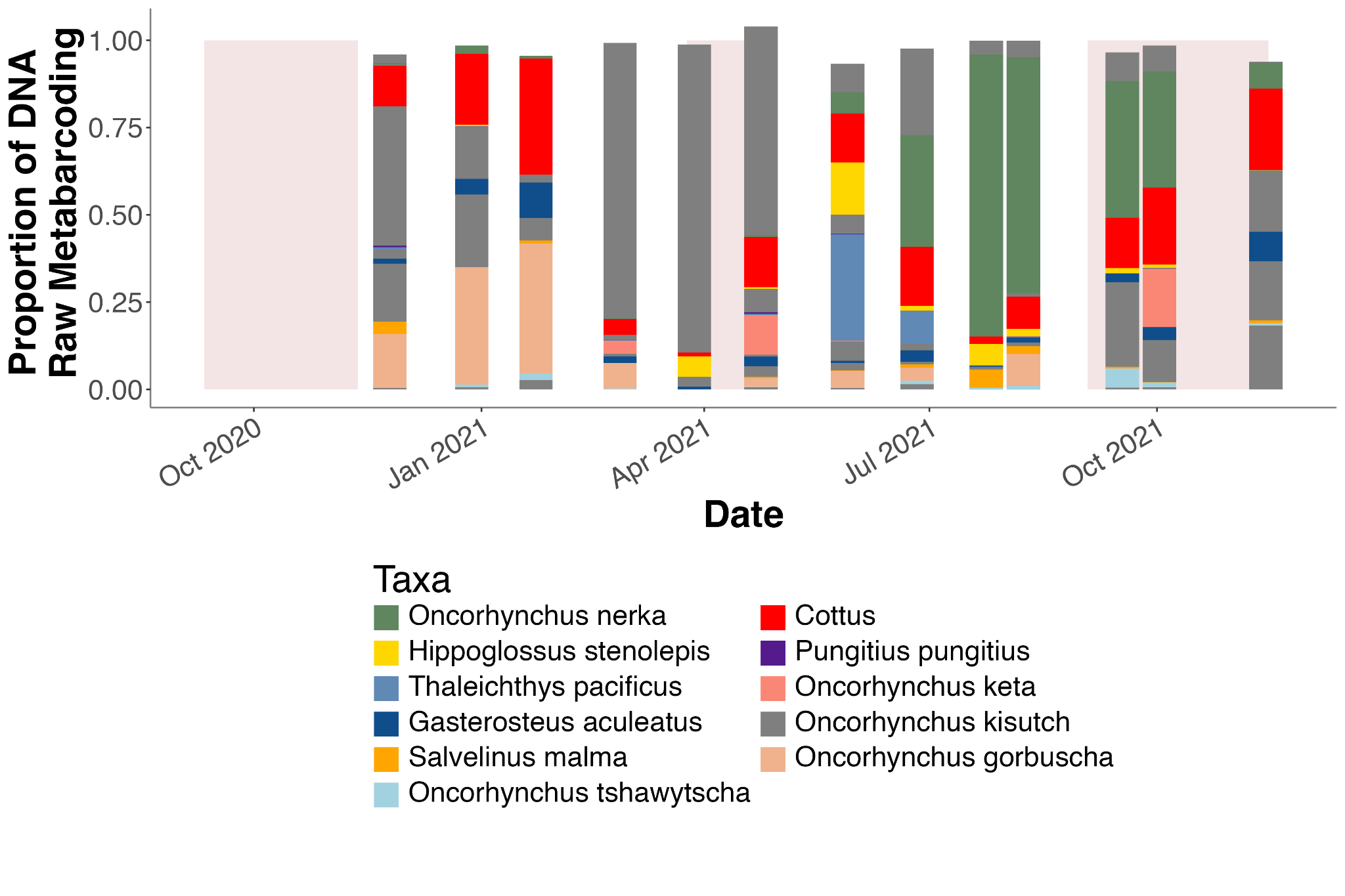

**Figure S6.7 Kenai River Raw Metabarcoding Results Over Time.**

Stacked bar plots of raw metabarcoding proportions. Taxa are color coded. Only bony fish species with at least 5 detections and 5,000 reads are shown. Periods of beluga occupancy are shaded in red.
